# Germ layer specific regulation of cell polarity and adhesion gives insight into the evolution of mesoderm

**DOI:** 10.1101/235267

**Authors:** Miguel Salinas-Saavedra, Amber Q. Rock, Mark Q Martindale

## Abstract

In triploblastic animals, Par-proteins regulate cell-polarity and adherens junctions of both ectodermal and endodermal epithelia. But, in embryos of the diploblastic cnidarian *Nematostella vectensis*, Par-proteins are degraded in all cells in the bifunctional gastrodermal epithelium. Using immunohistochemistry, CRISPR/Cas9 mutagenesis, and overexpression of specific mRNAs, we describe the functional association between Par-proteins, ß-catenin, and snail transcription factor genes in *N. vectensis* embryos. We demonstrate that the aPKC/Par complex regulates the localization of ß-catenin in the ectoderm by stabilizing its role in cell-adhesion, and that endomesodermal epithelial cells are organized by a different cell-adhesion system than that of overlying ectoderm. We also show that ectopic expression of *snail* genes, which are expressed in mesodermal derivatives in bilaterians, are sufficient to downregulate Par-proteins and translocate ß-catenin from the junctions to the cytoplasm in ectodermal cells. These data provide molecular insight into the evolution of epithelial structure and distinct mesodermal tissue in metazoan embryos.

## Introduction

Bilaterian animals comprise more than the 95% of the extant animals on earth and exhibit enormous body plan diversity (M. Q. Martindale & Lee, 2013). One of the most important morphological features in bilaterian evolution is the emergence of the mesoderm, an embryological tissue that gives rise important cell types such as muscle, blood, cartilage, bone, and kidneys in the space between ectoderm and endoderm. The emergence of mesoderm clearly contributed to the explosion of biological diversity throughout evolution (M. Martindale, 2005; M. Q. Martindale & Lee, 2013). Cnidarians (e.g., sea anemones, corals, hydroids, and “jellyfish”) are the sister group to bilaterians, and despite their surprisingly complex genomes (Putnam et al., 2007), do not possess a distinct mesodermal tissue layer. Instead, the gastrodermal lining to their gut cavity consists of a bifunctional endomesodermal epithelium with molecular characteristics of both bilaterian endodermal and myoepithelial mesodermal cells (Jahnel, Walzl, & Technau, 2014; M. Q. Martindale & Lee, 2013; M. Q. Martindale, Pang, & Finnerty, 2004; Technau & Scholz, 2003). For example, *brachyury* and *snail*, among other genes, contribute to the specification of the endomesodermal fates in both bilaterian and cnidarian embryos (Magie, Daly, & Martindale, 2007; M. Q. Martindale et al., 2004; Servetnick et al., 2017; Technau & Scholz, 2003; Yasuoka, Shinzato, & Satoh, 2016). Yet in bilaterians, mesodermal cells segregate from an embryonic endomesodermal precursor to form both endoderm and a third tissue layer (mesoderm) not present in the embryos of diploblastic cnidarians (Davidson et al., 2002; Maduro & Rothman, 2002; Q. Martindale et al., 2004; Rodaway & Patient, 2001; Solnica-Krezel & Sepich, 2012). How mesodermal cells originally segregated from an ancestral endomesodermal epithelium during animal evolution is still unclear (M. Martindale, 2005; M. Q. Martindale & Lee, 2013; Technau & Scholz, 2003), particularly because virtually all of the genes required for mesoderm formation are present in cnidarian genomes (Baumgarten et al., 2015; Chapman et al., 2010; Putnam et al., 2007; Shinzato et al., 2011). During the last decade, several studies have described molecular and cellular characteristics related to the segregation of mesoderm during bilaterian development (Darras, Gerhart, Terasaki, Kirschner, & Lowe, 2011; Keller, Davidson, & Shook, 2003; Schäfer, Narasimha, Vogelsang, & Leptin, 2014; Solnica-Krezel & Sepich, 2012). Here we investigate the cellular basis of morphogenesis during embryogenesis of the “diploblastic” sea anemone, *Nematostella vectensis*.

In most bilaterian embryos described to date, after a series of synchronous and stereotyped cleavage divisions, maternal determinants induce the localization of nuclear ß-catenin to blastomeres derived from the vegetal pole (M. Q. Martindale & Lee, 2013). Hence, gastrulation and the specification of endomesodermal fates is restricted to the vegetal pole. In these species, *brachyury* is expressed at the border of the blastopore and *snail* is expressed in the prospective mesodermal tissues (Technau & Scholz, 2003). The formation of mesoderm involves a variety of cellular processes including the downregulation of E-cadherin, loss of apicobasal cell polarity, and in some cases, the induction of epithelial-to-mesenchymal transition (EMT) (Acloque, Adams, Fishwick, Bronner-Fraser, & Nieto, 2009; Lim & Thiery, 2012; Schäfer et al., 2014; Solnica-Krezel & Sepich, 2012).

Embryos of the cnidarian starlet sea anemone *N. vectensis* develop without a stereotyped cleavage pattern but cell fates become organized along the embryonic animal-vegetal axis (Fritzenwanker, Genikhovich, Kraus, & Technau, 2007; Salinas-Saavedra, Stephenson, Dunn, & Martindale, 2015). During blastula formation, embryonic cells of *N. vectensis* form a single hollow epithelial layer. Epithelial cells of the animal pole, characterized by the nuclear localization of *Nv*ß-catenin prior to gastrulation (P. N. Lee, Kumburegama, Marlow, Martindale, & Wikramanayake, 2007; Wikramanayake et al., 2003), invaginate by apical constriction to form the endomesodermal epithelium (Magie et al., 2007; Tamulonis et al., 2011). The expression of *Nvbrachyury* around the presumptive border of the blastopore and *Nvsnail* genes in the presumptive endomesodermal gastrodermis of *N. vectensis* embryos occurs even before the morphological process of gastrulation begins (Röttinger, Dahlin, & Martindale, 2012; Scholz & Technau, 2003).

Interestingly, the components of the intracellular polarity Par system (*Nv*aPKC, *Nv*Par-6, *Nv*Par-3, *Nv*Par-1, and *Nv*Lgl), which show a highly polarized bilaterian-like subcellular distribution throughout all epithelial cells at the blastula stage in *N. vectensis* (Salinas-Saavedra et al., 2015), are specifically degraded and down-regulated from the endomesoderm during the gastrulation process (Figure 1A). We have previously suggested that the expression of bilaterian “mesodermal genes” (e.g. *Nvsnail)* might induce the loss of apicobasal cell-polarity indicated by the absence of the components of the Par system in the endomesoderm of *N. vectensis* embryos (Salinas-Saavedra et al., 2015). Recent studies in *N. vectensis* and bilaterians have provided information that supports this hypothesis. For example, it has been shown that *snail* is necessary and sufficient to downregulate Par3 in *Drosophila* mesoderm, inducing the disassembly of junctional complexes in these tissues (Weng & Wieschaus, 2016, 2017). In addition, we have shown that *Nvbrachyury* regulates epithelial apicobasal polarity of *N. vectensis* embryos, suggesting some aspects of epithelial cell polarity are highly conserved (Servetnick et al., 2017). Together, this evidence suggests a plausible cellular and molecular mechanism for the segregation of a distinct cell layer in bilaterian evolution from an ancestral bifunctional endomesodermal tissue. Thus, in this study we describe the functional association between the components of the Par system, apical junctions, epithelial integrity, and the nuclearization of *Nv*ß-catenin in a cnidarian embryo. In addition, we demonstrate that the endomesoderm in *N. vectensis* is organized by different junctional complexes that confer different functional properties to this tissue than the overlying ectoderm. And finally, we investigate the putative interactions between the components of the Par system, the canonical Wnt signaling pathway, and *snail* gene expression, giving insights on the evolution of the mesoderm and EMT.

**Figure 1.**
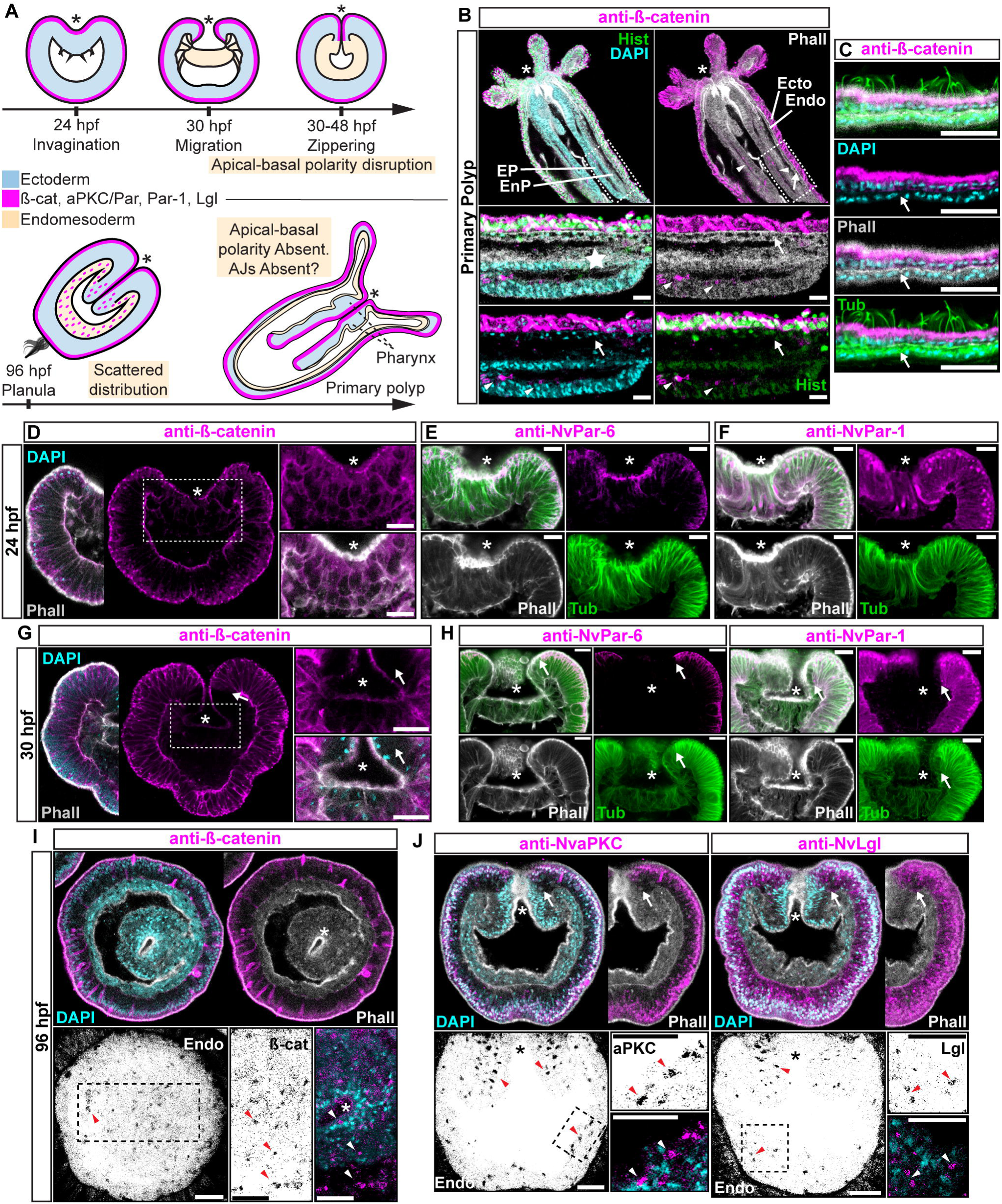
Components of the Par system and ß-catenin are downregulated from the *N. vectensis* endomesoderm during gastrulation. A-F. Confocal images of immunofluorescent staining (IFS) of lateral views of gastrulation embryos (animal pole up). The * marks the site of gastrulation in all cases. Samples are counterstained with Phalloidin (Phall) staining (white) to show cell boundaries, DAPI to visualize cell nuclei (blue), and Tubulin antibody (Tub) staining is shown as counterstain (green). All images are a single optical section from a z-stack confocal series. All scale bars, 20 µm. (A) Summary diagram depicting the localization of ß-catenin and Par proteins at the observed stages. Pale boxes denote changes observed in the endomesoderm. (B) IFS for ß-catenin (magenta) in primary polyps. High magnification images from boxed region (endomesoderm, Endo) are shown on the bottom. Arrows indicate the absence of ß-catenin expression in the endomesoderm. Arrowheads indicate the ß-catenin expression in the ectodermal pharynx (EP). Star indicates the endomesodermal pharynx (EnP). Histone antibody (Hist) staining is shown as counterstain to show the penetrability in the fixed tissue. See also Figure S1. (C) IFS for ß-catenin (magenta) in the ecto and endomesoderm (arrow) of primary polyps. (D) IFS for ß-catenin (magenta) at 24 hpf shows localization to the apical domain where adherens junctions reside in all cells of the blastula. High magnification images from boxed region (prospective endomesoderm) are shown on the right. (E) IFS for *Nv*Par-6 (magenta) at 24 hpf showing the same sub-cellular localization as ß-catenin (A). High magnification images from boxed region in (A) (prospective endomesoderm) are shown on the right. Merged image shown on upper left. (F) IFS for *Nv*Par-1 at 24 hpf shows a complementary basolateral expression. High magnification images from boxed region (prospective endomesoderm) are shown on the right. (G) IFS for ß-catenin at 30 hpf shows the loss of expression of ß-catenin (magenta) in invaginating endomesoderm (box). The arrow (D-F) marks the boundary between ectoderm and invaginating endomesoderm. High magnification images from boxed region (prospective endomesoderm) are shown on the right. (H) IFS for *Nv*Par-6 and *Nv*Par-1(magenta) at 30 hpf show that all Par proteins are down regulated at the site of gastrulation. IFS for *Nv*Par-6 shows an even earlier down regulation than ß-catenin (D). High magnification images from boxed region (prospective endomesoderm) are shown on the right. Merged image shown on upper left. (I) Oral view of IFS for ß-catenin (magenta) at 96 hpf showing apical localization in overlying ectoderm, but absence in endomesodermal tissues. The two bottom panels show high magnifications of the endomesoderm region (image inverted). Arrowheads indicate the localization of ß-catenin expression (black) in some scattered endomesodermal cells. (J) Lateral view of IFS for *Nv*aPKC and *Nv*Lgl (magenta) at 96 hpf showing loss of expression in invaginating epithelial cells. The four bottom panels show high magnifications of the endomesoderm region (image inverted). Arrowheads indicate the localization of *Nv*aPKC and *Nv*Lgl proteins (black) in some scattered endomesodermal cells.

## Results

### Ectodermal and endomesodermal epithelia are organized by different cell-cell adhesion complexes

Components of the Par system are not present in the cells of endomesodermal epithelium of *N. vectensis* during gastrulation, even though the very same cells express these proteins during the blastula stage (Salinas-Saavedra et al., 2015) (Figure 1). This absence is consistent with the absence of apical Adherens Junctions (AJs) in the endomesoderm of *N. vectensis* (Figure S1) and other cnidarians (Chapman et al., 2010; Ganot et al., 2015; Magie et al., 2007). At polyp stages, neither ß-catenin (an AJ-associated protein) (Figure 1B and 1C) nor the Par proteins (Figure S1C) are detectable in endomesodermal cells of either the gastrodermis or the pharynx. When *N. vectensis* embryos are stained with antibodies to ß-catenin (Figure 1) or if *Nv*ß-catenin::GFP mRNA is expressed in uncleaved zygotes (Figure S1B), clear localization of ß-catenin can be seen in the cortex of ectodermally derived epithelial cells (Figure 1B, 1C, 1D, 1G, and 1I), but not in endomesodermal cells (Figure 1B and 1C). In pharyngeal cells that are located between the epidermis and gastrodermis, *Nv*ß-catenin (Figure 1B and 1D), *Nv*Par-6 (Figure 1E), and *Nv*Par-1 (Figure 1F) expression begins to disappear, and is localized only in the most apical regions, indicating that AJs are being disassembled/degraded during the gastrulation process (Figure 1G and 1H). During later planula stages, ß-catenin and the components of the Par system display scattered patterns in the cytoplasm of a small subset of endomesodermal cells (Figure 1I and 1J). Even though we do not know the identity of these cells, this expression temporally coincides with the transient activation of Wnt signaling emanating from the oral pole (Kusserow et al., 2005; Marlow, Matus, & Martindale, 2013) at those developmental stages. In bilaterians (Acloque et al., 2009; Lim & Thiery, 2012) and *N. vectensis* (Kusserow et al., 2005; Marlow et al., 2013), the later activation of Wnt signaling is also associated with neurogenesis, and may cause the observed changes in protein localization.

Regardless of this scattered expression, it is clear that cells that undergo gastrulation in *N. vectensis* lose their polarized ectodermal cell-cell adhesion complex and components of the Par system, including ß-catenin, are downregulated from endomesodermal tissues (Figure 1). In bilaterians, the proper formation of an epithelial paracellular barrier (essential for tissue homeostasis) depends on the establishment of adhesive complexes between adjacent cells (Higashi, Arnold, Stephenson, Dinshaw, & Miller, 2016; Jonusaite, Donini, & Kelly, 2015), which are regulated by the aPKC/Par complex (Ohno, Goulas, & Hirose, 2015). To test if this absence of protein expression is correlated to differential cell-cell adhesion in the endomesodermal epithelium of *N. vectensis*, we assessed their role in regulating paracellular movements between ectodermal and endomesodermal epithelia by using a fluorescent tracer dye penetration assay (Figure 2A)(Higashi et al., 2016). For the purposes of these experiments, in order to avoid unwanted results related to tissue specification, cell proliferation, and cell movements, we used newly hatched juvenile polyps where the gastrodermis (endomesodermally derived) is fully differentiated.

**Figure 2.**
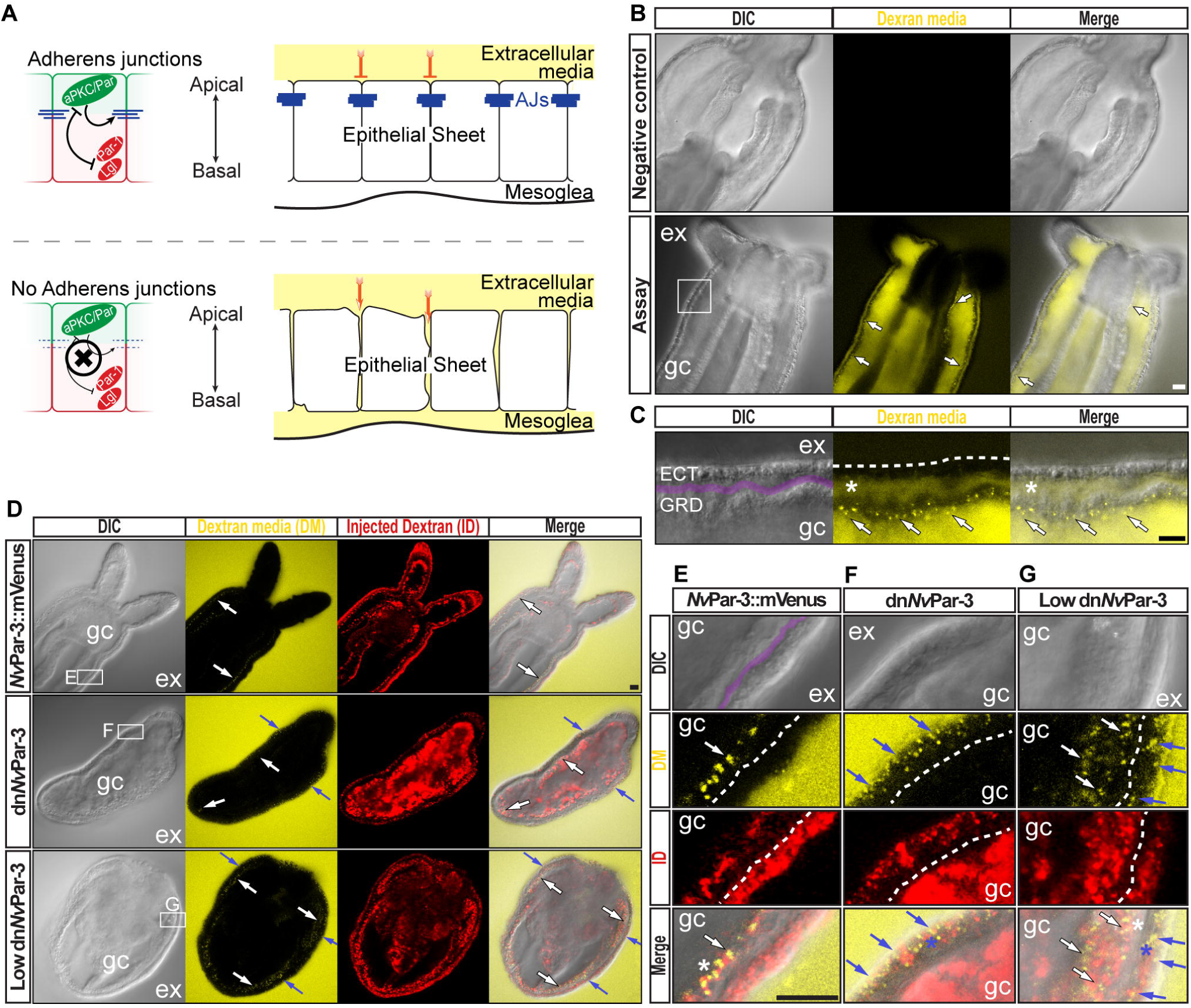
The aPKC/Par complex maintains Adherens Junctions (AJs) of ectodermal epithelial cells. Arrows indicate the direction of the flow: from gastric cavity (gc) to mesoglea (white) and from external media (ex) to ectoderm (blue). Dashed lines indicate the base of the epidermis. All images are single optical section from the z-stack confocal series. See also Figure S2 for dn*Nv*Par-3 description. Scale bars, 20 µm. (A) Diagram depicting the hypothesis that when the aPKC/Par complex is functional (top row), AJs are present (blue stripes) and a paracellular epithelial barrier is formed. When aPKC/Par complex is not functional (bottom row), AJs are disrupted, the epithelial barrier is perturbed, and the extracellular solution moves paracellularly into the mesoglea. (B) Penetration assay of wild type (uninjected) primary polyps at low magnification showing the movement of 10,000MW fluorescent dextran. Top row, no dextran. Bottom row, dextran (yellow) in the gc moves in to the mesoglea through paracellular spaces between gastrodermal cells (arrows). (C) High magnification images from box shown in (B). *: mesoglea (purple band) that separates the ectoderm (ECT, dashed line) from gastrodermis (GDR). Note the dye moving between cells from the gc media (arrows) (D) Low magnification images comparing polyps expressing *Nv*Par-3::mVenus and a dominant negative version of *Nv*Par-3 (dn*Nv*Par-3::mVenus) expressing-embryos. Dextran media (DM; extracellular) is pseudo-colored yellow. Dextran (red) was co-injected with mRNAs to label the cells and differentiate intracellular regions. mVenus channel was omitted for better visualization. Lower concentrations of dn*Nv*Par-3 were injected to preserve endomesodermal tissues. Note that the dextran media was found between the cells labeled in red. (E) High magnification images from (F) boxed region in (E). Purple band depicts Mesoglea. (F) High magnification images from (G) boxed region in (E). (G) High magnification images from (H) boxed region in (E). *: Paracellular spaces of both, the epidermis (blue) and gastrodermis (white).

*N. vectensis* polyps were exposed to media containing 10,000 MW fluorescent dextran (Molecular Probes, Inc.). When juvenile polyps are incubated in dextran for 5-10 minutes (Figure 2B), fluorescent dextran solution moves into the gastric cavity and then spreads into the mesoglea through the gastrodermal epithelium (Figure 2C) but does not enter the mesoglea through the outer ectodermally-derived epidermis (Figure 2C and 2D). These results suggest that cell-cell adhesion is differentially regulated between the epidermis and gastrodermis and the absence/disruption of AJs may compromise Septate Junctions (SJs) in the gastrodermis. Similar results were obtained in *N. vectensis* polyps when we overexpressed *Nv*Par-3::mVenus by injection of mRNA into uncleaved eggs which is normally expressed in ectodermal but not endodermal epithelial tissue (Figure 2D and 2E). However, in polyps expressing a dominant negative version of *Nv*Par-3::mVenus (dn*Nv*Par-3; microinjected into uncleaved eggs) dye penetrated between epithelial cells in both the gastrodermis and the outer epidermis (Figure 2D, 2F, and 2G), demonstrating an ancestral role of the aPKC/Par complex in the maintenance of cell-cell adhesion and the paracellular boundary (SJs) of epithelial cells during animal development.

### The *Nv*aPKC/Par complex regulates the formation and maintenance of cell-cell junctions

Our results suggest that the absence of Par proteins in the endomesoderm is associated with changes in cell-cell adhesion complexes. Pharmacological treatment of *N. vectensis* embryos with an aPKC activity inhibitor blocks cytokinesis but not mitosis in cleaving embryos (Figure 3A). In addition, a dominant negative version of *Nv*Par-1 (dn*Nv*Par-1), that lacks its kinase domain, localizes only to the cortex of cell-cell contacts (Figure 3B). dn*Nv*Par-1 can be phosphorylated by aPKC but cannot phosphorylate the aPKC/Par complex (Böhm, Brinkmann, Drab, Henske, & Kurzchalia, 1997; Vaccari, Rabouille, & Ephrussi, 2005). Thus, dn*Nv*Par-1 can localize to the cell cortex where aPKC is inactive. These results together suggest that the formation of cell-cell contacts is regulated by the activity of the aPKC/Par complex in *N. vectensis* embryos (Figure 3C). We further tested this hypothesis by using genome editing by CRISPR/Cas9 targeting *Nvpar-6* and *Nvpar-3* genes (Figure 3D). We did not observe any effects on the embryo until 36 hpf at 16°C (late blastula stage), indicating the activity of maternally loaded proteins up until that stage. When *Nv*Par-6 and *Nv*Par-3 are mutated, the ectodermal epithelium loses its integrity, presenting changes in thickness (Figure S2B and S3A), and interestingly, the endomesoderm (which does not express these proteins) generates cells with mesenchymal-like morphotypes that are never normally seen in this species (Figure 3D). In *Nvpar-6* and *Nvpar-3* mutant embryos, we also observed the disruption of microtubules and actin cytoskeleton (Figure S3B), and AJs (visualized with the ß-catenin antibody in Figure 3I) that confirms our previous observations of their role in regulating ectodermal cell polarity. Since we did not observe significant changes in the expression of germ layer markers when these genes were disrupted (e.g. *Nvbra*, *Nvsnail*, *NvSix3/6*, and *Nvfz10*; Figure S3E), we believe that only cell adhesion, but not cell specification, was affected by these experiments. Similar results were obtained when we overexpressed the mRNA encoding for a dominant negative version *Nv*Par-6 (dn*Nv*Par-6) and *Nv*Par-3 (dn*Nv*Par-3) into *N. vectensis* eggs (Figure S2 and S4). However, dominant negative effects on the injected embryos were observed at earlier stages (10-12 hpf) than the CRISPR/Cas9 mutants (zygotic expression) because the mutant proteins compete with the wild type proteins (maternally loaded). Hence, in these experiments, embryonic lethality (90%) and cell death were higher.

**Figure 3.**
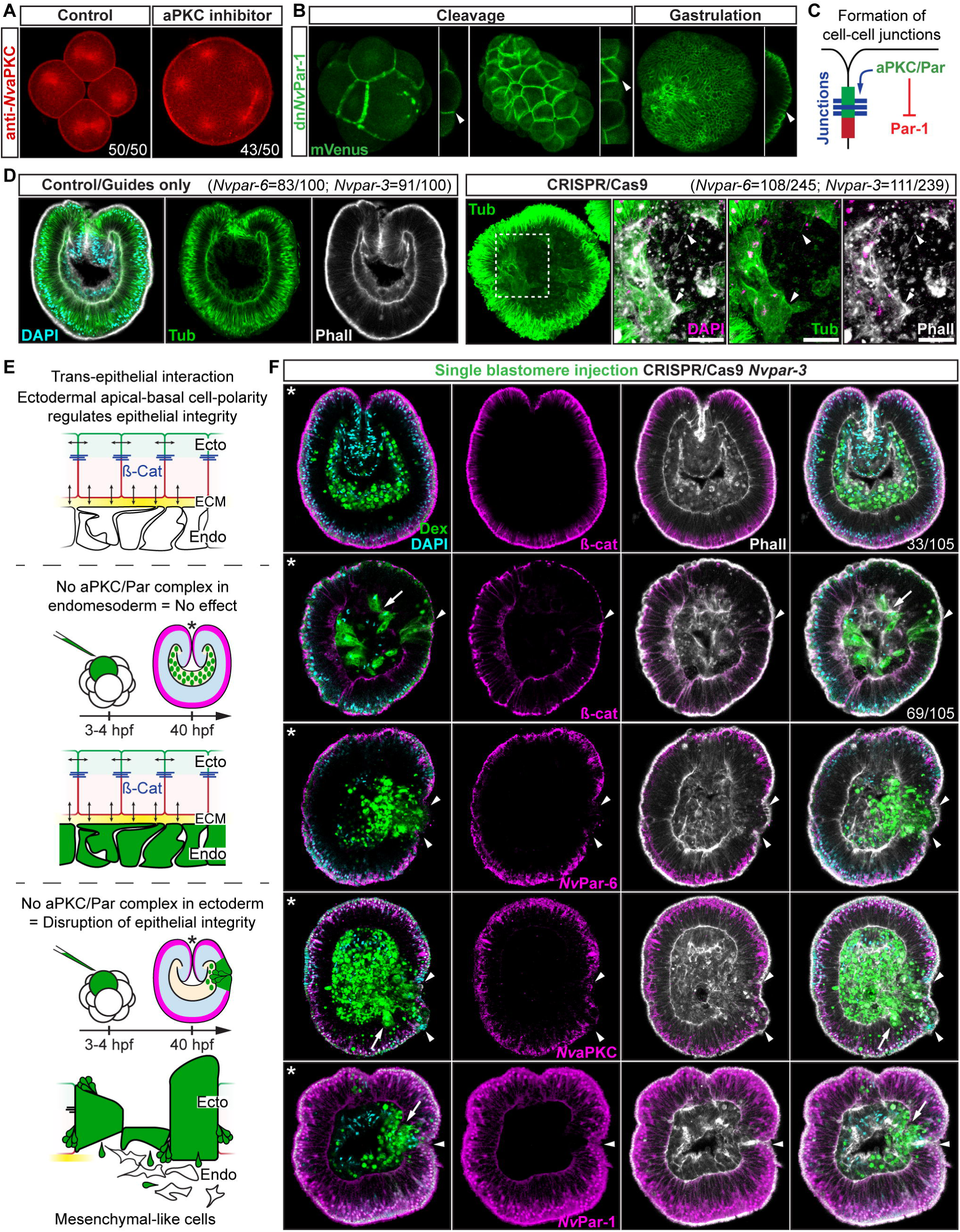
Ectodermal *Nv*aPKC/Par complex polarity regulates the epithelial integrity of both ecto- and endomesoderm. (A) IFS for *Nv*aPKC at 4 hpf showing that the aPKC inhibitor (Sigma P1614) blocks cytokinesis but not cell cycle. (B) In vivo expression of dn*Nv*Par-1 shows precocious localization to zones of cell contact during cleavage stages, well before wild-type *Nv*Par genes do. See also Figure S2A. (C) Diagram depicting the suggested the working hypothesis. (D) CRISPR/Cas9 knock-out for *Nv*Par-6 and *Nv*Par-3 at 48 hpf. Controls show no effect on gastrulation. Tubulin (Tub), Phalloidin (Phall), and DAPI are used as counterstains. CRISPR/Cas9 mutants: tubulin stained low magnification of CRISPR phenotype. High magnification images from boxed region shows mesenchymal-like cells. Arrowheads indicate filopodia-like structures. Number of cases observed for each gene are shown. See also Figure S2, S3, and S4. (E) Diagram depicts the hypothesis addressed in (I). The cell lineage derived from a single injected-blastomere is in green. (F) IFS for ß-catenin (ß-cat), *Nv*Par-6, *Nv*aPKC, and *Nv*Par-1 in single injected-blastomere CRISPR/Cas9 *Nv*Par-3 knock-outs at 40 hpf. Streptavidin-Biotin TxRed Dextran (Dex) is shown in green. Arrowheads indicate the absence of the protein and disrupted epithelium. Arrows indicate bottle-like shape cells. * indicate the orientation of the site of gastrulation. See also Figure S3D. Morphology is shown by DAPI, Tub, and Phall IFS. Except for 3B and 3D, all images are single optical sections from the z-stack confocal series. (B) and (D) are 3D reconstructions from a z-stack confocal series. All scale bars, 20 µm.

### The *Nv*aPKC/Par complex regulates transepithelial signaling

One surprising observation from the experiments described above show that the changes observed in the ectodermal and endodermal epithelium after disrupting *Nv*Par-6 and *Nv*Par-3 (Figure 3) suggests some sort of trans-epithelial regulation of cell-cell adhesion (most likely involving AJs) because these Par genes are not expressed in the endomesoderm. The polarizing activity of the aPKC/Par complex in the ectoderm is thus necessary to maintain the integrity of both ecto- and endodermal epithelia during cellular movements associated with gastrulation.

To assess whether the observed phenotypes on cell-cell adhesion are related to non-autonomous cell regulation (trans-epithelial interactions), we repeated the above experiments randomly injecting single blastomeres at 3-4 hpf (8-16 cell-stage) to make mutant clones in an otherwise wild type background. In these experiments, only the cell-lineage of the injected blastomere would be affected and would exhibit defective cell-cell adhesion in an otherwise undisturbed wild-type background. If endomesodermal cells derived from an injected blastomere display fibroblast/mesenchymal cell morphology, it would indicate that the organization of the endomesodermal epithelium is not dependent on the ectoderm but, rather, an intrinsic cell-autonomous activity of the aPKC/Par complex (Figure 3E). Our results show that only ectodermal- but not endomesodermal-lineages are affected by these mutations (Figure 3F and S3D). Presumptive ectodermal cells derived from an injected blastomere fail to maintain AJs (and potentially SJs) and the resulting clone of epithelial cells loses its structural integrity inducing cell extrusion. In contrast, presumptive endomesodermal cells derived from an injected blastomere develop into a normal endomesodermal epithelium (Figure 3F). Our results compliment the work of (Kirillova et al., 2018) and demonstrate that the proper cell-cell adhesion of the ectodermal layer somehow regulates trans-epithelially the integrity of the endomesodermal layer. This regulation may maintain the tension between cells during invagination at gastrula stages, or, in conjunction with the extracellular matrix (ECM) and basal cues, it may influence signaling patterns necessary to organize epithelial layers during *N. vectensis* embryogenesis.

### Interaction between the *Nv*aPKC/Par complex and the canonical Wnt signaling pathway

#### *Nv*aPKC/Par complex regulates ß-catenin localization

In bilaterians, AJs recruit members of the aPKC/Par complex and the direct interaction between Par-3 and aPKC/Par-6 is required for the maintenance and maturation of AJs (Ohno et al., 2015; Ragkousi, Marr, McKinney, Ellington, & Gibson, 2017). AJs are characterized by the binding between cadherins and ß-catenin: cadherins sequester ß-catenin from the cytoplasm to the cortex, making it unavailable for nuclear signaling and endomesoderm specification (Kumburegama, Wijesena, Xu, & Wikramanayake, 2011; Wikramanayake et al., 2003). Therefore, using ß-catenin as a marker for AJs, we separately co-injected *Nv*Par-3::mVenus or a mutated dn*Nv*Par-3::mVenus, with *Nv*ß-catenin::RFP into uncleaved zygotes. We observed cortical co-localization of *Nv*Par-3 and *Nv*ß-catenin at the cell boundaries in the ectodermal epithelium of embryos co-injected with *Nv*Par-3::mVenus and *Nv*ß-catenin::RFP (Figure 4A). However, in embryos co-injected with *Nv*ß-catenin::RFP and dn*Nv*Par-3::mVenus, we observed an alteration of the sub-cellular expression of *Nv*ß-catenin::RFP in all cells due to the translocation of ß-catenin from the cortical AJs into cell nuclei (Figure 4A).

**Figure 4.**
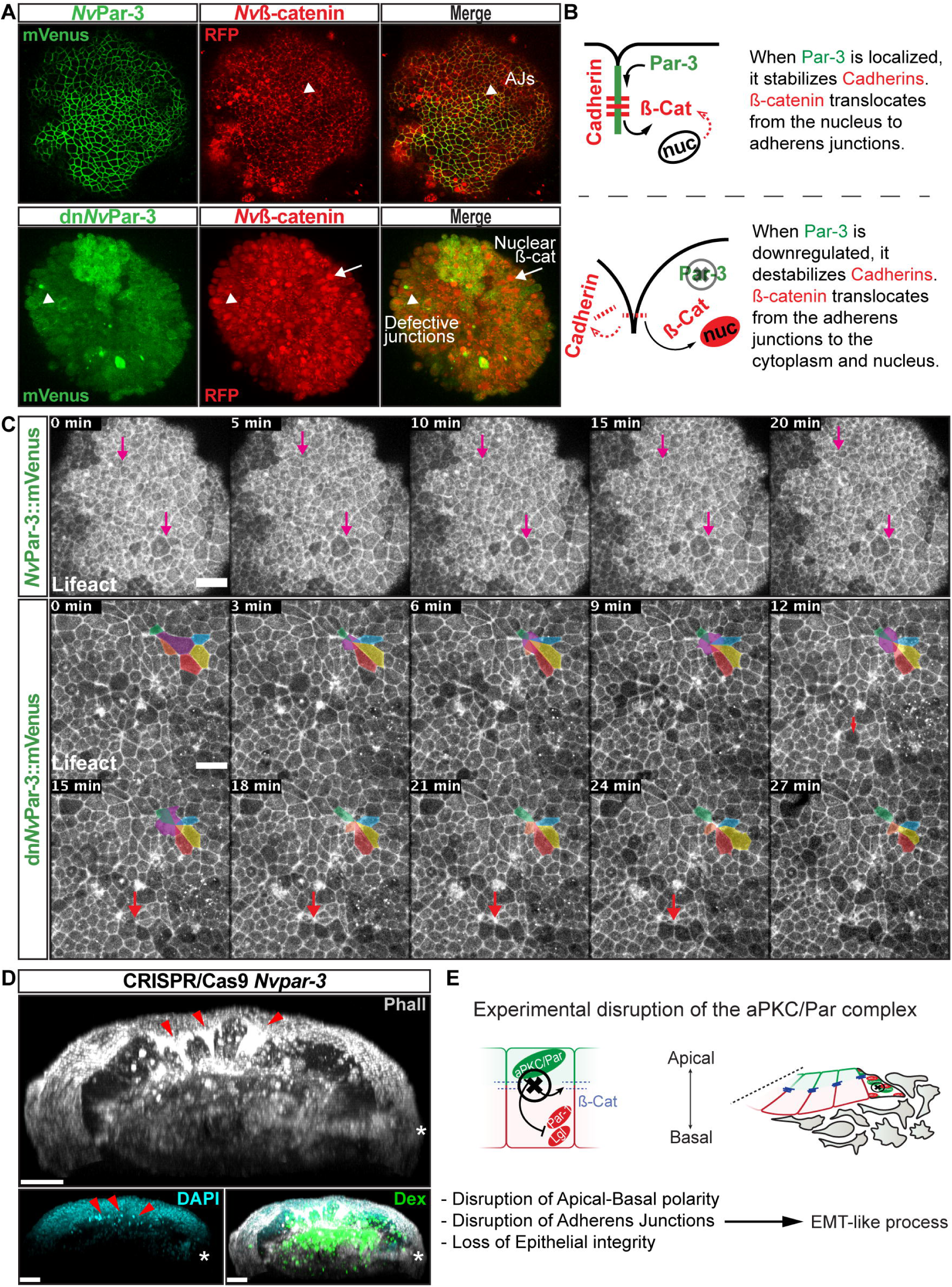
*Nv*aPKC/Par complex regulates ß-catenin localization and cell attachment. (A) In vivo co-localization of *Nv*Par-3venus co-injected with *Nv*ß-cateninRFP, and dn*Nv*Par-3venus co-injected with *Nv*ß-cateninRFP. Arrowheads indicate junctions (AJs). Arrows indicate nuclear ß-catenin. (B) Diagram of the suggested interpretation for A. (C) In vivo time series of ectodermal epithelial layers of embryos injected with *Nv*Par-3venus and dn*Nv*Par-3venus mRNA demonstrating epithelial delamination in the absence of functional *Nv*Par3. Lifeact::mTq2 mRNA was co-injected to visualize cell boundaries. Pink arrows indicate the absence cell detachments. A subset of cells was artificially colored. The purple cell detaches from the epithelium and the red arrow indicates a second cell detachment. See also Movie S1. (D) IFS of an embryo in which a single blastomere was injected with *Nv*Par-3 guide RNAs and Cas9 and green dextran. Red arrowheads indicate the apical constriction and delamination of ectodermal cells in the mutated clone of cells. Note the different layers of nuclei stained with DAPI. Asterisks indicate the site of gastrulation. (E) Diagram of the suggested interpretation for D. All images are 3D reconstructions from a z-stack confocal series. All scale bars, 20 µm.

In addition, results from *N. vectensis* embryos treated with 5µm 1-azakenpaullone (AZ; an inhibitor of GSK-3ß and a canonical Wnt agonist) suggest that GSK-3ß stabilizes AJs of epithelial cells in *N. vectensis* embryos (Figure S5). We observed an expansion of the expression domain of Par-6 (Figure S5) and a stabilization of AJs (labeled with ß-catenin in Figure S5) in endomesodermal cells of treated embryos, which was never observed in control embryos.

Interestingly, the association between the nuclearization of ß-catenin (canonical Wnt signaling pathway) and the Par system has been poorly studied. Two studies, one in *Drosophila* (Sun et al., 2001) and the another in *Xenopus* (Ossipova, Dhawan, Sokol, & Green, 2005) embryos, have shown by immunoblotting that the kinase Par-1 is associated with Dishevelled protein and might act as a positive regulator of Wnt signaling. Here, we show *in vivo* embryonic evidence suggesting that *Nv*Par-3 (whose cortical localization is normally inhibited by Par-1) recruits *Nv*ß-catenin protein and stabilizes its localization at the apico-lateral cortex of ectodermal cells through the formation of AJs. Furthermore, the putative disassembly of the aPKC/Par complex induced by dn*Nv*Par-3 overexpression, induces the nuclearization of *Nv*ß-catenin protein (Figure 4A) due to its cytosolic availability caused by AJs disruption. Strikingly, we also observed the extrusion of individual cells from the ectodermal epithelium of dn*Nv*Par-3 treated-embryos (Figure 4C) and single injected-blastomere CRISPR/Cas9 *Nv*Par-3 knock-out (Figure 4D). This suggests that these treatments induce EMT-like processes, not observed under control conditions (Figure 4C).

Thus, our data suggest that preexisting mechanisms downstream to the induction of EMT may have been redeployed to segregate layers during the evolution to bilaterians. Bringing the question whether or not endomesodermal genes would induce similar effects when they are expressed in *N. vectensis* embryos.

We have recently showed that *Nvbrachyury* regulates apicobasal polarity of epithelial cells in *N. vectensis* embryos (Servetnick et al., 2017). We, therefore, examined the role of *Nvsnail* genes on the localization of ß-catenin, components of the Par system, and the stabilization of AJs. Our hypothesis is that expression of *N. vectensis snail* genes would destabilize AJs and induce the nuclearization of ß-catenin in ectodermal epithelial cells.

### *Nvsnail* genes induce the translocation of *Nv*ß-catenin from AJs to the cytoplasm

*N. vectensis* has two *snail* genes, *Nvsnail-A* and *Nvsnail-B*, which are both expressed in the endomesodermal plate prior to and throughout the gastrulation process, and which define the boundary between gastrodermis and ectodermal pharynx (Amiel et al., 2017; Magie et al., 2007; Röttinger et al., 2012). To determine the role of *Nvsnail* genes on ß-catenin nuclearization, we co-injected the mRNA of *Nv*Snail-A::mCherry, *Nv*Snail-B::mCherry, and *Nv*ß-catenin::GFP into uncleaved eggs. The overexpression of both proteins *Nv*Snail-A::mCherry and *Nv*Snail-B::mCherry together induce the ectopic translocation of *Nv*ß-catenin::GFP to the nuclei of ectodermal cells (Figure 5A). This treatment also delocalizes *Nv*Par-3 from the cell cortex when both *Nv*Snail::mCherry proteins are co-expressed with *Nv*Par-3::mVenus (Figure 5B).

**Figure 5.**
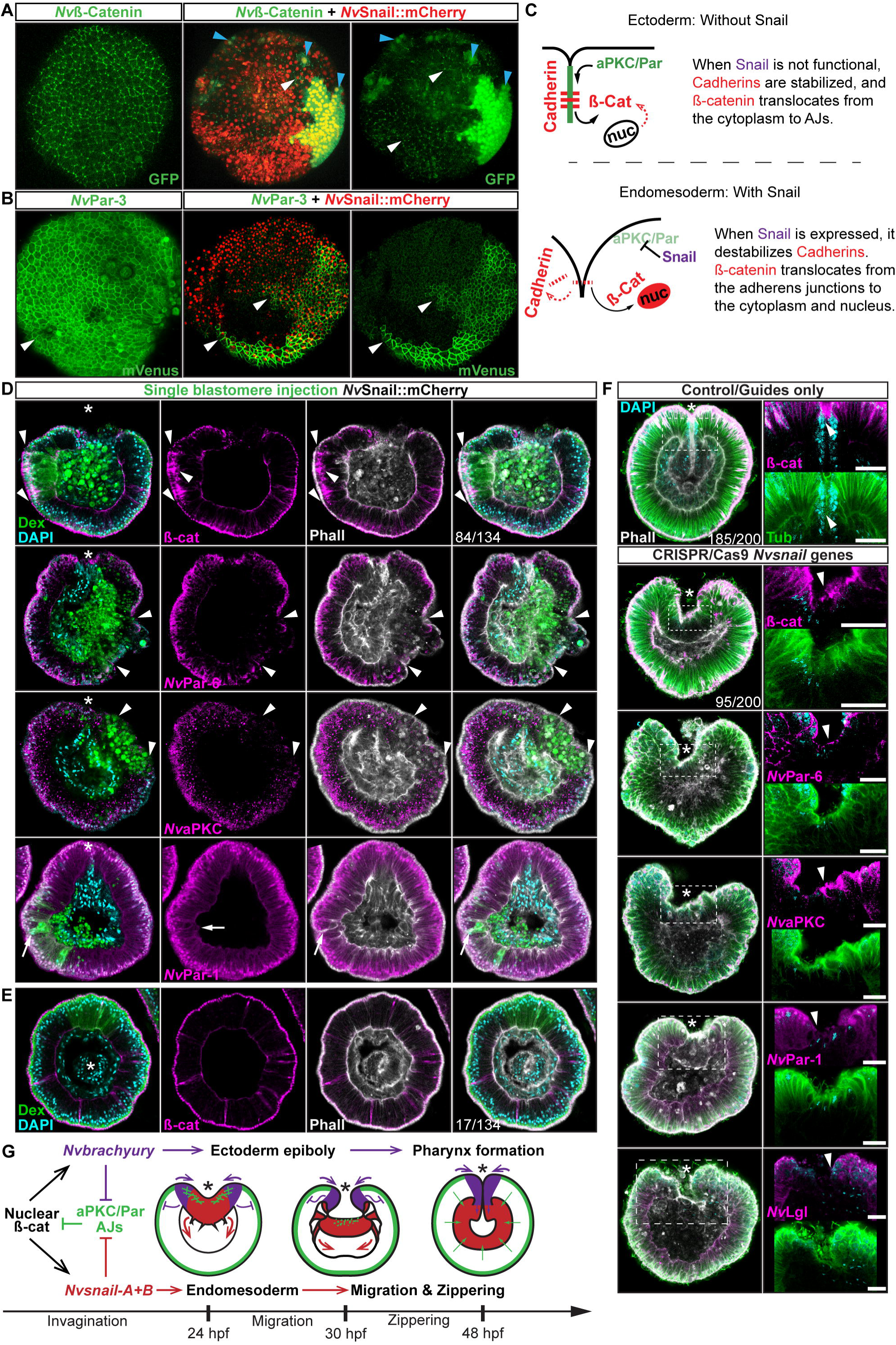
*Nvsnail* genes downregulate AJs and *Nv*aPKC/Par polarity allowing endomesodermal migration. (A) in vivo localization of *Nv*ß-cateninGFP co-injected with both *Nv*Snail-A::mCherry and *Nv*Snail-B::mCherry mRNA together in zygotes at 40 hpf. White arrowheads indicate AJs. Patched patterns of cytosolic and nuclear ß-catenin (Blue arrowheads) were observed. (B) In vivo localization of *Nv*Par-3::mVenus co-injected with both *Nv*Snail-A::mCherry and *Nv*Snail-B::mCherry mRNA together at 40 hpf. Patched patterns of AJs (White arrowheads) were observed. (C) Diagram depicts the suggested interpretation for A and B. (D) IFS for ß-catenin (ß-cat), *Nv*Par-6, *Nv*aPKC, and *Nv*Par-1 in embryos at 40 hpf where *Nv*Snail-A::mCherry and *Nv*Snail-B::mCherry mRNA were overexpressed together into a single ectodermal blastomere lineage (followed by green Streptavidin-TxRed Dextran (Dex). Arrowheads indicate the absence of the protein, cytosolic ß-cat, and disrupted epithelium. Arrows indicate bottle-like shape cells. *site of gastrulation. (E) IFS for ß-cat in embryos at 40 hpf where *Nv*Snail-A::mCherry and *Nv*Snail-B::mCherry mRNA were overexpressed together into a single blastomere lineage and no affects were observed. See also Figure S7. (F) Embryo wide CRISPR/Cas9 knock-out for both *Nvsnail-A* and *Nvsnail-B* at 40 hpf showing that AJs form in presumptive endomesodermal region similar to ectodermal cells. High magnification images from boxed region (endomesodermal plate) are shown on the right. Arrowheads indicate protein localization. (G) Graphical summary of the observed results with previous published data (Servetnick et al., 2017). Morphology is shown by DAPI, Tub, and Phall IFS. Except from 6A and 6B, all images are single optical sections from the z-stack confocal series. (A) and (B) are 3D reconstructions from a z-stack confocal series. All scale bars, 20 µm.

To determine the role of *Nvsnail* genes on cell adhesion/epithelial polarity, we randomly injected single blastomeres at the 8-32 cell-stage with mRNA from both *Nv*Snail-A::mCherry and *Nv*Snail-B::mCherry together. The fluorescent dextran that was co-injected with the mRNAs could be used to detect the clones where the over-expression of the co-injected mRNAs occurred in a “wild-type” background (Figure 5D). Similar to the *Nvpar-3* knock-out (Figure 3F), the expression of *Nvsnail* genes is sufficient to induce the degradation of Par proteins and AJs (ß-catenin) from the ectoderm and disrupts its epithelial integrity; however, nuclear ß-catenin was not observed under these treatments (Figure 5D). Thus, nuclear *Nv*ß-catenin::GFP observed *in vivo* when we overexpressed *Nv*Snail proteins (Figure 5A) is a consequence of the high cytosolic availability generated by its ectopic overexpression and release from AJs.

Interestingly, not every ectodermal cell was affected by these treatments even though all of the cells expressed the injected mRNAs (Figure 5A, 5E, and S6). This patched pattern suggests that the response to *Nvsnail* over-expression is spatially regulated. These results suggest that the role of *Nvsnail* genes on AJs and apicobasal cell polarity is constrained to the site of gastrulation in *N. vectensis* embryos under natural conditions, and that these genes may be required for gastrulation movements. Therefore, we predicted that ß-catenin (AJs) and Par proteins will be retained in the cells of the *N. vectensis* endomesodermal plate if both *Nvsnail* genes are disrupted.

### *Nvsnail* genes downregulate apicobasal cell polarity and AJs in the endomesodermal epithelium of *N. vectensis* embryos

The *snail* genes temporally down-regulate E-cadherin during mesoderm segregation and EMT in bilaterian animals (Lim & Thiery, 2012). As we have shown here, as well as in previous studies (Magie et al., 2007; Magie & Martindale, 2008), the cells comprising the endomesodermal plate lose their cell-cell adhesion during gastrulation in *N. vectensis* embryos. It may be possible that temporal regulation of endomesodermal patterning might act upon the AJs. Our data suggest that once gastrulation is complete and the pharynx forms, components of the Par system and the ß-catenin components of the AJs are degraded from both the cortex and cytoplasm of endomesodermal cells (Figure 1 and S1). Hence, it could be possible that *Nvbrachyury* induces the disruption of apicobasal polarity (Servetnick et al., 2017), remnant AJs maintain the endomesodermal-plate cells together, and *Nvsnail* genes degrades and prevents the reassembly of AJs in the endomesoderm of *N. vectensis*.

To address these issues, we used CRISPR/Cas9 knock-out of *Nvsnail-A* and *Nvsnail-B* genes together to inhibit zygotic function of these genes and investigate their role on the temporal regulation of AJs and cell polarity. In CRISPR/Cas9 mutants, the endomesodermal plate forms but it does not migrate further than its first invagination during gastrulation (Figure 5F). Furthermore, AJs (labeled with ß-catenin) and apical Par proteins (labeled with anti*Nv*Par-6 and anti*Nv*aPKC) are retained at the apical cortex of the cells of the endomesodermal plate (Figure 5F and S6). Surprisingly, *Nv*Par-1 and *Nv*Lgl were not detected in those cells (Figure 5F), suggesting that the degradation of these basolateral proteins precede or do not depend on the activity of the *Nvsnail* genes. This suggests that *Nvsnail* regulates apical cell-polarity, AJs turnover, and the migration (‘zippering’) but not the invagination of the endomesodermal plate during gastrulation of *N. vectensis* embryos. Interestingly, the invagination of the endomesodermal plate (controlled by the Wnt/PCP pathway) is uncoupled from its specification in *N. vectensis* embryos (Kumburegama et al., 2011; N. M. Wijesena, 2012), which is consistent with our observations.

## Discussion

### AJs are down-regulated in mesoderm and neural crest of bilaterian animals

The segregation of different germ layers during embryogenesis of many bilaterian animals is carried out by similar cellular mechanisms. EMT is a shared mechanism utilized by mesoderm, neural crest cell (NCC), and tumorigenesis to delaminate cells in bilaterian animals (triploblastic animals). During EMT, the nuclearization of ß-catenin induces the expression of ‘endomesodermal’ genes like *brachyury* and *snail* (Acloque et al., 2009). The expression of these genes downregulates epithelial cadherins, disrupts apicobasal polarity (mediated by the aPKC/Par complex), disassembles AJs, and induces changes in cytoskeleton organization (Acloque et al., 2009; Lim & Thiery, 2012). A rearrangement of the actin-myosin cytoskeleton induces apical constriction of cells (generating a bottle-like shape), which detach from the epithelial sheet, break down the basal membrane, and invade a specific tissue as mesenchymal cells (Acloque et al., 2009; Lim & Thiery, 2012; Ohsawa, Vaughen, & Igaki, 2018).

Interestingly, mesoderm formation, tumorigenesis, and EMT have never been described as natural processes during *N. vectensis* (a diploblastic animal) embryogenesis. During *N. vectensis* gastrulation (Magie et al., 2007; Tamulonis et al., 2011), cells around the edge of the blastopore at the animal pole (which expresses *Nvbrachyury*) acquire a bottle-like shape by apical constriction, leading to epithelial buckling and the invagination of presumptive endomesoderm (which expresses *Nvsnail*). However, throughout this process the endomesoderm remains as a monolayer of epithelial cells and individual mesenchymal cells never detach and invade the blastocoel. We have shown that by disrupting the aPKC/Par complex (apicobasal cell-polarity) in *N. vectensis* (Figure 3 and 4), we are able to convert cells from the endomesodermal epithelium into mesenchymal-like cells, translocate *Nv*ß-catenin (Figure 4A), and emulate EMT-like processes (apical constriction and individual cell-detachments) in the ectodermal epithelium of *N. vectensis* treated-embryos (Figure 4C and 4D). These results demonstrate that the cnidarian *N. vectensis* possesses mechanisms necessary to segregate individual germ layers (e.g. mesoderm and NCC) described in bilaterians; however, they do not do it.

Our working hypothesis is that the *N. vectensis* embryo is composed of two independent morphogenetic modules that are integrated and organized by the pharynx (Figure 5G). The first observation is that the ectoderm, whose apicobasal polarity (and thus, AJs and epithelial integrity) is regulated by *Nvbrachyury* that promotes ectodermal epithelial morphogenesis and pharynx formation (Servetnick et al., 2017), and the second module is generated by endomesodermal differentiation and cell-movements that are regulated by *Nvsnail* genes. This is supported by the expression *Nvbrachyury* (Servetnick et al., 2017)^9^ in *Nvsnail* knock-out embryos (Figure S7), and *Nvsnail* knock-out phenotypes where ectodermal pharynx develops normally but no clear endomesoderm is formed (Figure S6). Additional work is required to elucidate any differences in function between *Nvsnail-A* and *Nvsnail-B* genes, however, both modules are specified by nuclear ß-catenin (Röttinger et al., 2012), suggesting that the nuclear ß-catenin (maternal) shift from the animal pole in cnidarians to the vegetal pole in bilaterians is mechanistically plausible and sufficient to re-specify the site of gastrulation and germ-layers along the animal-vegetal axis during Metazoan evolution (P. N. Lee et al., 2007; M. Q. Martindale & Lee, 2013).

### The dual identity and collective migration of the endomesodermal cells

Bilaterian-EMT has been a focus of study for decades as a mechanism to segregate different cell layers involved in a variety of different normal and pathological biological processes (Nieto, Huang, Jackson, & Thiery, 2016; Ohsawa et al., 2018). This process appears to depend on the fine regulation of *snail* expression levels and their temporal activity. For example, during NCC migration, cells display ‘partial-EMT’ where cells remain attached to several neighboring cells but their apicobasal polarity and AJs are down-regulated, allowing collective-cell migration (J. M. Lee, Dedhar, Kalluri, & Thompson, 2006; Nieto et al., 2016; Ribeiro & Paredes, 2014; Theveneau & Mayor, 2013; Weng & Wieschaus, 2017). Our data suggest that ‘partial-EMT’ may be the mechanism by which the endomesodermal epithelium migrates into the blastocoel in *N. vectensis* embryos during normal gastrulation (Figure 6). In this scenario, upstream factors that regulate *snail* transcription may be critical for this process.

**Figure 6.**
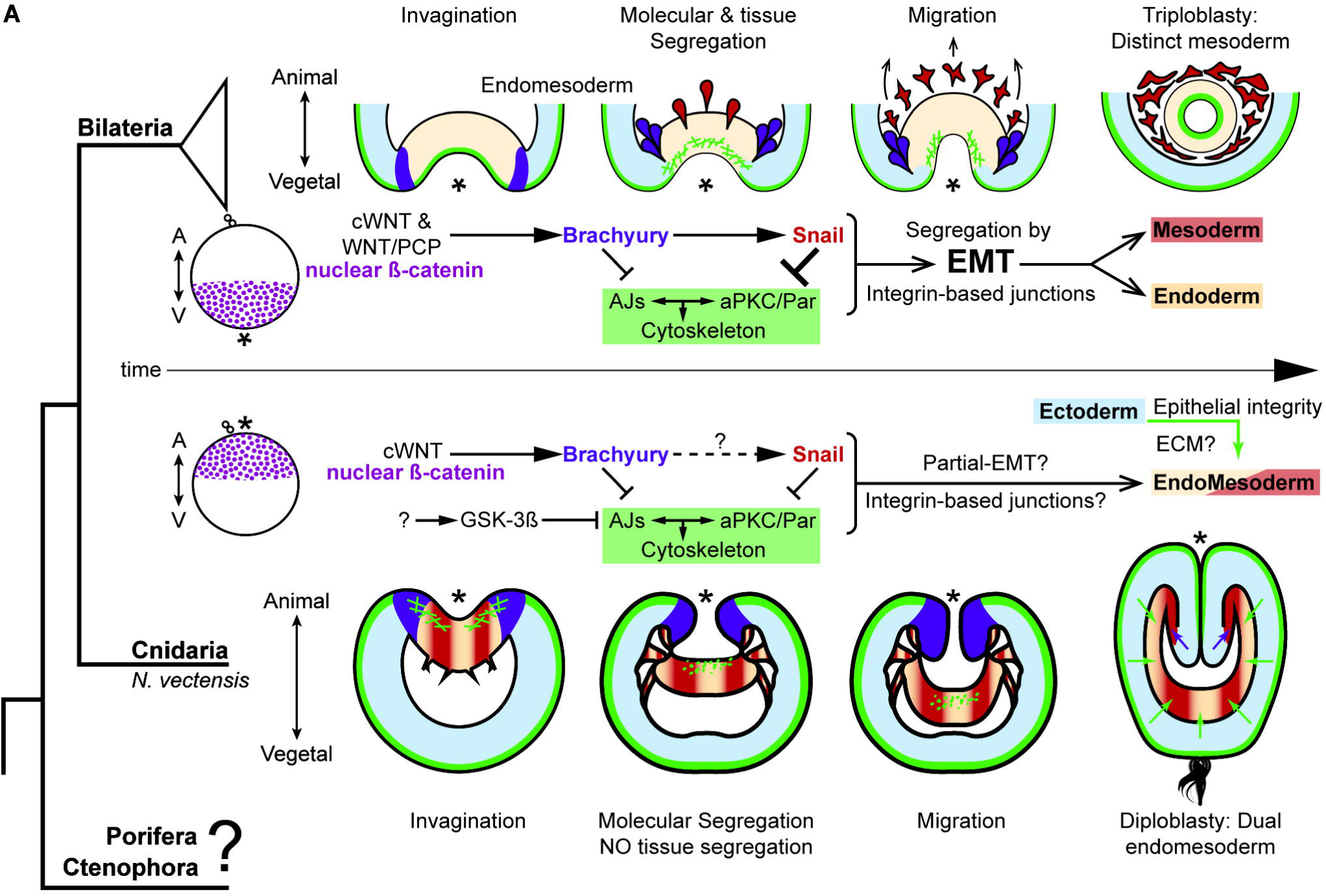
The differences between epithelial structure in ectoderm and endomesoderm in *N. vectensis* embryos are due to the lack of mechanisms to segregate a distinct mesoderm. (A) Diagram depicting key cellular and molecular mechanisms involved during gastrulation of bilaterian and *N. vectensis* (a cnidarian) embryos. See also Figure S8.

In bilaterian animals, there are many other pathways in addition to the canonical Wnt pathway that activate *snail* transcripton and induce the disruption of AJs and apicobasal cell polarity. For example, TGFß, BMP, NANOS, FGF, and MEK/ERK/ERG take on roles during the specification of mesoderm, NCC migration, tumorigenesis, and other EMT-related processes (Barrallo-Gimeno & Nieto, 2005; Lim & Thiery, 2012; Nieto et al., 2016). Concordantly in *N. vectensis* embryos, cells of the pharyngeal and endomesodermal tissues express components of all these pathways (Amiel et al., 2017; Extavour, Pang, Matus, & Martindale, 2005; Matus, Thomsen, & Martindale, 2006, 2007; Röttinger et al., 2012; N. Wijesena, Simmons, & Martindale, 2017) that may modify their cellular characteristics. For example, one cadherin (*Nv*CDH2 (Clarke, Miller, Lowe, Weis, & Nelson, 2016): 1g244010), and kinases that modify tubulin and histones are differentially regulated between ecto- and endomesodermal epithelium (N. Wijesena et al., 2017).

In conclusion, *N. vectensis* has both up and downstream cellular and molecular mechanisms associated with EMT described in bilaterians. However, *N. vectensis* does not segregate a distinct mesodermal germ layer nor display EMT under natural conditions. In bilaterians, this mechanism must have evolved to segregate mesodermal cells from the endoderm to retain the tight cell-cell junctions required in endodermal epithelia. Interestingly, mesoderm segregation via EMT in *Drosophila* takes place after epithelial folding in response to *snail* expression. In these embryos, contractile myosin enhances the localization of AJs and Par-3 in the presumptive mesoderm and prevents their downregulation by Snail, thus delaying EMT (Weng & Wieschaus, 2016, 2017). Furthermore, the overexpression of Snail in *Drosophila* embryos is sufficient to disassemble ectodermal-AJs, but mesodermal-AJs are maintained by actomyosin contraction that antagonize Snail effects (Weng & Wieschaus, 2016, 2017). Our results suggest a similar mechanism since *Nvsnail* overexpression in endomesodermal lineages (Figure S6) is not sufficient to segregate cells and the endomesoderm remains as an epithelium. However, unlike *Drosophila*, Par proteins and AJs are not enhanced but degraded during the gastrulation of *N. vectensis* (Figure 1). As it is discussed in (Weng & Wieschaus, 2017), not only the degradation but also the turnover of AJs and Par proteins in adjacent epithelia is essential for EMT-mediated germ layer segregation in different animals. The dual identity of *N. vectensis* endomesoderm is characterized by the continuous expression of *Nvsnail* genes (M. Q. Martindale et al., 2004) that repress the turnover of AJs and may play a role in inhibiting EMT from occuring (Figure 5).

Interestingly, components of the Wnt/PCP pathway are expressed only in the endomesoderm (Kumburegama et al., 2011; N. M. Wijesena, 2012), while components of the Par system are expressed only in the ectoderm (Salinas-Saavedra et al., 2015). It could be that NvSnail degrades AJs and inhibits their re-assembly in the endomesoderm, but the activation of contractile myosin by the Wnt/PCP pathway maintains the endomesodermal cells together in *N. vectensis* embryos. Hence in bilaterians, a mechanism (most likely downstream of Snail) that connects the cytoskeleton with cell-polarity may have evolved to tighten cell-cell adhesion in the endoderm and allow EMT.

To elucidate this, further comparative research and funding are needed to understand the cellular mechanisms that evolve to segregate mesoderm and control epithelial cell polarity at the base of the metazoan tree. For example, ctenophores segregate a mesodermal cell population during embryogenesis but do not have the genes that encode for all cell-cell adhesion complexes and specify for mesoderm in bilaterians (Figure S8) (Ganot et al., 2015; Ryan et al., 2013). Thus, there is much to be learned by the comparative study of cell biology.

## Material and methods

### Culture and spawning of *Nematostella vectensis*

Spawning, gamete preparation, fertilization and embryo culturing of *N. vectensis* (RRID:SCR_005153) embryos was performed as previously described (Hand & Uhlinger, 1992; Layden, Röttinger, Wolenski, Gilmore, & Martindale, 2013; Röttinger et al., 2012; Wolenski, Layden, Martindale, Gilmore, & Finnerty, 2013). Adult *N. vectensis* were cultivated at the Whitney Laboratory for Marine Bioscience of the University of Florida (USA). Males and females were kept in separate glass bowls (250 ml) in 1/3x seawater (salinity: 12pp) reared in dark at 16°C. Animals were fed freshly hatched *Artemia* 3 times a week and macerated oyster the day before spawning. Spawning was induced by incubating the adults under an eight-hour light cycle at 25°C the night before the experiment. Distinct groups of animals were spawned once every 2 weeks. Oocytes and sperm were collected separately and fertilized *in vitro* by adding sperm to egg masses for 25 minutes. The jelly mass surrounding the fertilized eggs was removed by incubating the eggs in 4% L-Cysteine (in 1/3x seawater; pH 7.4) for 15-17 minutes and then washed 3 times with 1/3x seawater. De-jellied eggs were kept in glass dishes (to prevent sticking) in filtered 1/3 seawater at 16°C until the desired stage.

### Immunohistochemistry

All immunohistochemistry experiments were carried out using the previous protocol for *N. vectensis* (Salinas-Saavedra et al., 2015) with a slight modification in the glutaraldehyde concentration to allow better antibody penetration. Embryos were fixed on a rocking platform at room temperature in two consecutive steps. Embryos of different stages were fixed for no longer than 3 minutes in fresh Fix-1 (100mM HEPES pH 6.9; 0.05M EGTA; 5mM MgSO4; 200mM NaCl; 1x PBS; 3.7% Formaldehyde; <u>0.2%</u> Glutaraldehyde; 0.5% Triton X-100; and pure water). Then, Fix-1 was removed and replace with fresh Fix-2 (100mM HEPES pH 6.9; 0.05M EGTA; 5mM MgSO4; 200mM NaCl; 1x PBS; 3.7% Formaldehyde; <u>0.05%</u> Glutaraldehyde; 0.5% Triton X-100; and pure water). Embryos were incubated in Fix-2 for 1 hour. Fixed embryos were rinsed at least five times in PBT (PBS buffer plus 0.1% BSA and 0.2% Triton X-100) for a total period of 3 hours. PBT was replaced with 5% normal goat serum (NGS; diluted in PBT) and fixed embryos were blocked for 1 to 2 hours at room temperature with gentle rocking. Primary antibodies were diluted in 5% NGS to desired concentration. Blocking solution was removed and replaced with primary antibodies diluted in NGS. All antibodies incubations were conducted over night on a rocker at 4°C. After incubation of the primary antibodies, samples were washed at least five times with PBT for a total period of 3 hours. Secondary antibodies were then applied (1:250 in 5% NGS) and samples were left on a rocker overnight at 4°C. Samples were then washed with PBT and left on a rocker at room temperature for an hour. To visualize F-actin, samples were incubated then for 1.5 hours in Phalloidin (Invitrogen, Inc. Cat. # A12379) diluted 1:200 in PBT. Samples were then washed once with PBT and incubated with DAPI (0.1µg/µl in PBT; Invitrogen, Inc. Cat. # D1306) for 1 hour to allow nuclear visualization. Stained samples were rinsed again in PBS two times and dehydrated quickly into isopropanol using the gradient 50%, 75%, 90%, and 100%, and then mounted in Murray’s mounting media (MMM; 1:2 benzyl benzoate:benzyl alcohol) for visualization. Note that MMM may wash DAPI out of your sample. For single blastomere microinjection experiments, after Phalloidin staining, samples were incubated with Texas Red Streptavidin (1:200 in PBT from 1mg/ml stock solution; Vector labs, Inc. Cat.# SA-5006. RRID:AB_2336754) for 1 hour to visualize the injected dextran. We scored more than 1,000 embryos per each antibody staining and confocal imaged more than 50 embryos at each stage.

The primary antibodies and concentrations used were: mouse anti-alpha tubulin (1:500; Sigma-Aldrich, Inc. Cat.# T9026. RRID:AB_477593), rabbit anti-ß-catenin (1:300; Sigma-Aldrich, Inc. Cat.# C2206. RRID:AB_476831), mouse anti-histone H1 (1:300; F152.C25.WJJ, Millipore, Inc. RRID:AB_10845941). Rabbit anti-*Nv*aPKC, rabbit anti-*Nv*Lgl, rabbit anti-*Nv*Par-1, and rabbit anti-*Nv*Par-6 antibodies are custom made high affinity-purified peptide antibodies that were previously raised by the same company (Bethyl Inc.). All these four antibodies are specific to *N. vectensis* proteins (Salinas-Saavedra et al., 2015) and were diluted 1:100.

Secondary antibodies are listed in Supplementary file 1.

### Fluorescent tracer dye penetration assay

Primary polyps were incubated and mounted in 1/3 sea water with fluorescent dextran solution (0.5 mg/ml). For uninjected embryos we used Dextran, Alexa Fluor® 555 (Molecular Probes, INC. Cat.# D34679). For injected embryos, expressing fluorescent proteins, we used Dextran, Alexa Fluor® 647 (Molecular Probes, INC. Cat.# D22914). Animals were observed within 10 minutes of incubation. 15 animals were recorded per treatment. For better visualization of the dextran solution inside the gastric cavity as shown in Figure 2B, we delivered additional dextran solution by microinjecting dye through the polyp’s mouth. For the rest of the experiments, we mainly focused in the ectodermal permeability and we let the polyps to eat the solution by themselves as grown babies.

### mRNA microinjections

The coding region for each gene of interest was PCR-amplified and cloned into pSPE3-mVenus or pSPE3-mCherry using the Gateway system (Roure et al., 2007). Eggs were injected directly after fertilization as previously described (Layden et al., 2013; Salinas-Saavedra et al., 2015) with the mRNA encoding one or more proteins fused in frame with reporter fluorescent protein (N-terminal tag) using final concentrations of 450 ng/µl for each gene. Fluorescent dextran was also co-injected to visualize the embryos. For single blastomere microinjections, we raised the embryos until 8-16 cell stages (3-4 hpf) and co-injected the mRNA solution with Biotinylated Dextran Amine-Texas Red (10 µg/µl; Vector labs, Inc. Cat.# SP-1140. RRID:AB_2336249). Live embryos were kept at 16°C and visualized after the mRNA of the FP was translated into protein (2-3 hours). To avoid lethality, lower mRNA concentrations of the mutant proteins (250 ng/µl) were used to image the specimens for Figures 2 and 4, and Movie S1. Live embryos were mounted in 1/3 sea water for visualization. Images were documented at different stages from 3-96 hrs. post fertilization. We injected and recorded more than 500 embryos for each injected protein and confocal imaged approximately 20 specimens for each stage for detailed analysis of phenotypes *in vivo*. We repeated each experiment at least five times obtaining similar results for each case. The fluorescent dextran and primers for the cloned genes are listed in Supplementary file 1.

### CRISPR/Cas9 knock-outs

To target our gene of interest, we used synthetic guide RNAs (sgRNA; Synthego, Inc.) and followed the instructions obtained from the manufacturer to form the RNP complex with Cas9 (Cas9 plus sgRNAs). Target sites, off-target sites, and CFD scores were identified and sgRNA were designed using CRISPRscan (Doench et al., 2014; Moreno-Mateos et al., 2015). We delivered the RNP complex by microinjection as previously described (Ikmi, McKinney, Delventhal, & Gibson, 2014; Servetnick et al., 2017; N. Wijesena et al., 2017). Lyophilized Cas9 (PNA Bio., Inc. Cat.# CP01) was reconstituted in nuclease-free water with 20% glycerol to a final concentration of 2µg/µl. Reconstituted Cas9 was aliquoted for single use and stored at -80°C. Embryos were injected, as described for mRNA microinjections, with a mixture (12.5µl) containing sgRNAs (80 ng/μl of each sgRNA), Cas9 (3 μg), and Alexa Fluor 488-dextran (0.2 μg/μl; Molecular Probes, Inc. Cat.# D22910). Cas9 and sgRNA guides only controls were injected alongside each round of experiments. sgRNA guides controls are only shown in figures as Cas9 had no significative effects. 3 sgRNA were used to knock out *Nvpar-3*, 3 sgRNA were used to knock out *Nvpar-6*, 6 sgRNA were used to knock out *Nvsnail-A*, and 6 sgRNA were used to knock out *Nvsnail-B*. Single-embryo genomic DNA was analyzed as previously described (Servetnick et al., 2017). Gene expression was confirmed by i*n situ* hybridization. We injected and recorded more than 1000 embryos for each treatment. We repeated each experiment at least six times obtaining similar results for each case. sgRNAs’ sequences and PCR primers flanking the targeted region are listed in Supplementary file 1.

### *In situ* hybridization

*In situ* hybridization was carried out following a previously published protocol for *N. vectensis* (Wolenski et al., 2013). Animals were fixed in ice-cold 4% paraformaldehyde with 0.2% glutaraldehyde in 1/3x seawater for 2 min, followed by 4% paraformaldehyde in PBTw for 1 hour at 4°C. Digoxigenin (DIG)-labeled probes, previously described (Röttinger et al., 2012; Salinas-Saavedra et al., 2015), were hybridized at 63°C for 2 days and developed with the enzymatic reaction of NBT/BCIP as substrate for the alkaline phosphatase conjugated anti-DIG antibody (Roche, Inc. Cat.#11093274910. RRID:AB_514497). Samples were developed until gene expression was visible as a purple precipitate.

### Drug treatment

We incubated *N. vectensis* embryos in 20µM of aPKC pseudosubstrate inhibitor (Protein kinase Cζ pseudosubstrate, myristoyl trifluoroacetate salt, Sigma, Cat.#P1614) from 0 to 4 hpf. Controls and 1-azakenpaullone (AZ; Sigma, Cat.#A3734) drug treatment of *N. vectensis* embryos was performed as previously described (Leclère, Bause, Sinigaglia, Steger, & Rentzsch, 2016; Röttinger et al., 2012). Embryos were developed in 5µm AZ from 3 to 76 hpf. Controls were incubated in 0.08% DMSO.

### Imaging of *N. vectensis* embryos

Images of live and fixed embryos were taken using a confocal Zeiss LSM 710 microscope using a Zeiss C-Apochromat 40x water immersion objective (N.A. 1.20). Pinhole settings varied between 1.2-1.4 A.U. according to the experiment. The same settings were used for each individual experiment to compare control and experimental conditions. Results from *in situ* hybridization studies were imaged using a Zeiss Imager.M2 with a Zeiss 425 HRc color digital camera run by Zeiss Zen 2012 software. Z-stack images were processed using Imaris 7.6.4 (Bitplane Inc.) software for three-dimensional reconstructions and FIJI for single slice and movies. Final figures were assembled using Adobe Illustrator and Adobe Photoshop.

## Acknowledgments

We thank Leslie S. Babonis for TEM data in Figure S1, A. Wikramanayake for *Nv*ß-catenin expression constructs, and C. Magie and E. Röttinger for *Nvsnail-A* expression constructs. We thank E. Seaver, C.E. Schnitzler, and the members of Martindale’s lab for helpful discussion. We also thank the *NSF-IOS 1239422- Broadening participation of underrepresented groups in Developmental Biology”* and NSF REU (DBI-1156528) programs for AQR. This research was supported by the NIH GM093116, NASA 16-EXO16_2-0041, NSF IOS-1755364, and Synthego grants to MQM.

## Author Contributions

MS-S., and MQM. designed research and analyzed data. MS-S. performed research with help of AQR and MQM. MS-S., and MQM. wrote the manuscript with help of AQR. All authors read and approved the final manuscript.

## Declaration of Interests

The authors declare no competing interests.

## Supplemental Information

**Figure S1. Related to Figure 1. Epidermal and gastrodermal cells are joined by different set of junctional complexes.**

*(A)* TEM micrographs of *N. vectensis* primary polyps. Epidermal cells (ectodermally derived) are joined most likely by AJs (Red arrowheads). Gastrodermal cells (endomesodermally derived) are interconnected by fewer and shorter contacts, most likely by septate junctions (yellow arrowheads). a and b: two different types of epidermal cells. Note the ectodermally-derived cnidocyte in b. c and d: two different types of gastrodermal cells. b’ and d’ are high magnification images from boxed region in b and d, respectively. gc: gastric cavity. Scale bars in a, b, c, and d: 500 nm. Scale bars in b’ and d’: 200 nm.

*(B) In vivo* localization of *Nv*ß-catenin::GFP after 36 hpf in *N. vectensis* embryos. Arrowheads indicate the cortical localization of *Nv*ß-catenin::GFP (AJs) in the ectoderm that was not observed in the endomesoderm. High magnification image from boxed region is shown on the right. Scale bars: 20 µm.

*(C)* Immunofluorescent staining for *Nv*Par-6 and *Nv*Par-1(red) at late planula and polyp stages show that both Par proteins are absented from the endomesoderm. Phalloidin is shown in gray. Histone and tubulin antibody staining are shown in Figure 1B and 1C as counterstain to show the penetrability in the fixed tissue. Scale bars: 20 µm.

**Figure S2. Related to Figure 2 and 3. Disruption of the aPKC/Par complex in *N. vectensis* embryos.**

(A) Diagram depicting the modifications made to *Nv*Par-1, *Nv*Par-6, and *Nv*Par-3 sequence to generate the dominant negative version of each protein (dn*Nv*Par-1, dn*Nv*Par-6, and dn*Nv*Par-3, respectively), which lack the putative interaction domain with *Nv*aPKC. In the right, a diagram depicts the localization of *Nv*Par-1, *Nv*aPKC, *Nv*Par-6, and *Nv*Par-3 proteins in epithelial cells. The putative interaction with *Nv*aPKC, restricts the localization of *Nv*Par-6 and *Nv*Par-3 strictly to the apical cortex of the cell, and *Nv*Par-1 to the lateral cortex of the cell. See also Figure S3.

(B) Immunofluorescent staining for Tubulin and Phalloidin at gastrula stage of embryos expressing dn*Nv*Par-6, and dn*Nv*Par-3. The overexpression of either dn*Nv*Par-6::mVenus or dn*Nv*Par-3::mVenus induced phenotypes where the endomesodermal cells (yellow arrows) are disorganized during gastrulation. We observed a disorganized endoderm formed by 1) cells with fibroblast-like morphologies, 2) stellate shaped mesenchymal-like cells, or 3) a mass of round dead cells. Cell morphotypes are outlined in red and their schematic representation is presented below them. The penetrance of the obtained phenotypes when either dn*Nv*Par-6::mVenus (blue) or dn*Nv*Par-3::mVenus (magenta) are overexpressed is indicated for each case.

(C) Immunofluorescent staining for Tubulin and Phalloidin at polyp stage of embryos expressing dn*Nv*Par-6, and dn*Nv*Par-3. We observed ‘Endoderm-less’ polyps from phenotype 3: absence of an organized endoderm, tentacles or complete mesenteries in injected animals that survived to this stage (2 weeks post fertilization). A recognizable ectodermal pharynx (yellow arrowheads) was detected in some of these polyps.

(D) The position of the sgRNAs (red) and primers (green) used for the PCR assay are shown on the diagram depicting the genomic sequence of *Nvpar-6*. Note the absence of fragments of *Nvpar-6* (arrow) resulting from CRISPR/Cas9 mediated mutagenesis. The presence of other bands suggests mosaicism. Black rectangles correspond to *Nvpar-6* exon. PF: primer forward. PF: primer reverse.

(E) The position of the sgRNAs (red) and primers (green) used for the PCR assay are shown on the diagram depicting the genomic sequence of *Nvpar-3*. Note the absence/truncation of fragments of *Nvpar-3* (arrow) resulting from CRISPR/Cas9 mediated mutagenesis. The presence of other bands suggests mosaicism as shown in Figure S3E. Blue rectangles correspond to *Nvpar-3* exon. PF: primer forward. PF: primer reverse.

**Figure S3. Related to Figure 3. CRISPR/Cas9 mediated mutagenesis of *Nvpar-3.*** Morphology is shown by DAPI, Tubulin, and Phalloidin staining. *: site of gastrulation is up.

(A) Different ectodermal thickness (yellow) observed in CRISPR/Cas9 mutants. Control and affected (thick and thin) epithelia were aligned at the base of the ectoderm (red line) for better visualization.

(B) Immunofluorescent staining for cytoskeleton and *Nv*Par-6 in CRISPR/Cas9 *Nvpar-6* knock-out embryos (mosaic phenotype). Arrowheads indicate the absence of *Nv*Par-6. High magnification images are shown on the right. The cytoskeleton is apically organized only where *Nv*Par-6 is apically localized.

(C) Diagram depicts the role of aPKC/Par complex (green) on cytoskeleton.

(D) Immunofluorescent staining for ß-catenin (ß-cat) in single injected-blastomere control and CRISPR/Cas9 *Nvpar-3* knock-outs at 40 hpf. Streptavidin-Biotin TxRed Dextran (Dex) is shown in green. Arrowheads indicate the absence of the protein and disrupted epithelium. Arrows indicate bottle-like shape cells. Note the displacement of nuclei (yellow arrow) and changes in tubulin (Tub) staining.

(E) *In situ* hybridization of *Nvpar-3* knockout embryos (Cas9 and gRNAs) compared with control embryos at 40 hpf. The disruption of the *N. vectensis* Par/aPKC complex modified the morphology but did not modify the cell-fate specification of endomesodermal cells. Lower panel for *Nvfz-10* may represent an extreme case of mesenchymal-like endomesoderm.

**Figure S4. Related to Figure 3 and S2. *In vivo* localization of dn*Nv*Par-6 and dn*Nv*Par-3 proteins at different embryonic stages.**

Images of the whole embryo correspond to a 3D reconstruction from a z-stack series. Side panels are a single optical section from the z-stack series. An aboral view is shown for all gastrula stages. Yellow arrows indicate the apico-lateral cortex labeled with Lifeact::mTq2. Scale bars: 10µm.

(A) *In vivo* localization of *Nv*Par-6::mVenus and dn*Nv*Par-6::mVenus at cleavage, blastula, and gastrula stages. *Nv*Par-6::mVenus distributes uniformly at the apical region of the cell but displays a scattered pattern. However, dn*Nv*Par-6::mVenus displays stronger cortical localization due to its interaction with *Nv*Cdc42 at the apical and apico-lateral cortex of the cells. This was confirmed by their co-expression with *Nv*Cdc42::mCherry. White arrowheads indicate the scattered localization of *Nv*Par-6::mVenus. White arrows indicate the cortical and stronger localization of dn*Nv*Par-6::mVenus.

(B) *In vivo* localization of Lifeact::mTq2 in *Nv*Par-6::mVenus and dn*Nv*Par-6::mVenus expressing embryos shown in (A) at cleavage, blastula, and gastrula stages. The actin cytoskeleton was also affected by the overexpression of dn*Nv*Par-6::mVenus.

(C) *In vivo* co-distribution of *Nv*Cdc42::mCherry with *Nv*Par-6::mVenus at blastula and gastrula stages.

(D) I*n vivo* co-distribution of *Nv*Cdc42::mCherry with dn*Nv*Par-6::mVenus at blastula and gastrula stages. *Nv*Cdc42::mCherry localization is also affected when dn*Nv*Par-6::mVenus is overexpressed.

(E) Graphical summary of the observed results for *Nv*Par-6::mVenus and dn*Nv*Par-6::mVenus.

(F) *In vivo* localization of *Nv*Par-3::mVenus and dn*Nv*Par-3::mVenus at cleavage, blastula, and gastrula stages. dn*Nv*Par-3 displays broader localization, resembling the localization of *Nv*Par-6 and indicating its release from AJs (compare with *Nv*Par-3). White arrowheads indicate the punctate localization of *Nv*Par-3::mVenus. White arrows indicate the broader localization of dn*Nv*Par-3::mVenus.

(G) *In vivo* localization of Lifeact::mTq2 in *Nv*Par-3::mVenus and dn*Nv*Par-3::mVenus expressing embryos shown in (F) at cleavage, blastula, and gastrula stages. The actin cytoskeleton was also affected by the overexpression of dn*Nv*Par-3::mVenus.

(H) *In vivo* co-distribution of *Nv*Cdc42::mCherry with *Nv*Par-3::mVenus at blastula and gastrula stages.

(I) I*n vivo* co-distribution of *Nv*Cdc42::mCherry with dn*Nv*Par-3::mVenus at blastula and gastrula stages. *Nv*Cdc42::mCherry localization is not affected when dn*Nv*Par-3::mVenus is overexpressed.

(J) Graphical summary of the observed results for *Nv*Par-3::mVenus and dn*Nv*Par-3::mVenus.

**Figure S5. Apical junctions are regulated by GSK-3ß in epithelial cells during gastrulation in *N. vectensis* embryos.**

(A) IFS for *Nv*Par-6 in AZ-treated embryos.

(B) IFS for ß-catenin in AZ-treated embryos.

(C) IFS for *Nv*Par-1 in AZ-treated embryos.

(D) Graphical summary of the observed results.

Par-6, ß-catenin, and Par-1 were detected at the apical cortex (arrowheads) of the endomesodermal epithelium of AZ-treated embryos, but not in control embryos. Arrows indicated the stabilization of microtubules at the apical cortex of endomesodermal cells. Two distinct phenotypes were observed after AZ treatment: (a’) gastrulation without elongation 27% (27/100), (b’) no invagination of the endomesoderm 13% (13/100).

**Figure S6. Related to Figure 5. *Nvsnail-A and Nvsnail-B* together regulate AJs.**

*site of gastrulation.

(A) Immunofluorescent staining of embryos at 40 hpf where *Nv*Snail-A::mCherry and *Nv*Snail-B::mCherry mRNA were overexpressed together into a single blastomere lineage (followed by green Streptavidin-TxRed Dextran (Dex). No affects were observed in approximately 1/8 of the injected embryos. Arrowheads indicate the disrupted epithelium. Arrows indicate bottle-like shape cells.

(B) Embryo wide CRISPR/Cas9 knock-out for both *Nvsnail-A* and *Nvsnail-B* at 40 hpf showing embryos that did not gastrulate and retain ß-catenin (ß-cat) and *Nv*aPKC at the apical cell-cortex of the endomesodermal plate (arrowheads).

(C) Embryo wide CRISPR/Cas9 knock-out for both *Nvsnail-A* and *Nvsnail-B* at 40 hpf showing embryos that developed a pharynx (arrow) but not an organized endomesoderm.

(D) Embryo wide CRISPR/Cas9 knock-out for both *Nvsnail-A* and *Nvsnail-B* at 2 weeks hpf showing embryos that did not developed a gastrodermal epithelium.

**Figure S7. Related to Figure 5. CRISPR/Cas9 mediated mutagenesis of *Nvsnail-A and Nvsnail-B*.**

(A) The position of the sgRNAs (red) and primers (green) used for the PCR assay are shown on the diagram depicting the genomic sequence of *Nvsnail-A*. Note the absence of fragments of *Nvsnail-A* (arrow) resulting from CRISPR/Cas9 mediated mutagenesis. The presence of other bands suggests mosaicism. Black rectangles correspond to *Nvsnail-A* exon. Blue double arrow depicts UTR regions. PF: primer forward. PF: primer reverse.

(B) The position of the sgRNAs (red) and primers (green) used for the PCR assay are shown on the diagram depicting the genomic sequence of *Nvsnail-B*. Note the absence of fragments of *Nvsnail-B* (arrow) resulting from CRISPR/Cas9 mediated mutagenesis. The presence of other bands suggests mosaicism as shown in (C). Black rectangles correspond to *Nvsnail-B* exon. Blue double arrow depicts UTR regions. PF: primer forward. PF: primer reverse.

(C) *In situ* hybridization of *Nvsnail-A+Nvsnail-B* knockout embryos (Cas9 and gRNAs) compared with control embryos at 40 hpf. *Nvsnail-A+Nvsnail-B* knockout embryos display no expression and mosaic expression of *Nvsnail-A* and *Nvsnail-B* but did not modify the cell-fate specification of ectodermal cells.

**Figure S8. Related to Figure 6. Suggested model for mesoderm specification in Metazoa.**

(A) Differential regulation of cell adhesion by the endomesoderm GRN is mediated by changes in cell polarity that regulate ß-catenin localization. These mechanisms emerged at the Bilateria + Cnidaria node.

(B) Ctenophores do not possess a full complement of cell adhesion, cell-polarity, and endomesodermal GRN components present in the most common ancestor between Cnidaria and Bilateria. However, ctenophores possess a distinct mesoderm, suggesting the emergence of different mechanisms to segregate mesoderm in Metazoa.

**Movie S1. Related to Figure 4. EMT-like process occurs in dn*Nv*Par-3::mVenus expressing cells.** Lifeact::mTq2 is shown in grey. Arrows indicate the apical constriction of detaching cells. Scale bars:10µm

