## Supplementary Materials for "Germ layer specific regulation of cell polarity and adhesion gives insight into the evolution of mesoderm"

**Supplementary file 1**

| REAGENT or RESOURCE | SOURCE | IDENTIFIER |
| --- | --- | --- |
| **Antibodies** | | |
| Mouse Anti-alpha-Tubulin Monoclonal Antibody, Unconjugated, Clone DM1A | Sigma-Aldrich | T9026; RRID:AB_477593 |
| Anti-beta-Catenin antibody produced in rabbit | Sigma-Aldrich | C2206; RRID:AB_476831 |
| Anti-Histone, H1 + core proteins, clone F152.C25.WJJ antibody | Millipore | MABE71; RRID:AB_10845941 |
| anti-NvaPKC custom peptide antibody produced in rabbit | Bethyl labs; Salinas-Saavedra et al., 2015 |  |
| anti-NvLgl custom peptide antibody produced in rabbit | Bethyl labs; Salinas-Saavedra et al., 2015 |  |
| anti-NvPar-1 custom peptide antibody produced in rabbit | Bethyl labs; Salinas-Saavedra et al., 2015 |  |
| anti-NvPar-6 custom peptide antibody produced in rabbit | Bethyl labs; Salinas-Saavedra et al., 2015 |  |
| DAPI (4',6-Diamidino-2-Phenylindole, Dihydrochloride) | Thermo Fisher Scientific | D1306; RRID:AB_2629482 |
| Alexa Fluor 488 Phalloidin | Thermo Fisher Scientific | A12379; RRID:AB_2315147 |
| Goat anti-Mouse IgG Secondary Antibody, Alexa Fluor 568 | Thermo Fisher Scientific | A-11004; RRID:AB_2534072 |
| Goat anti-Rabbit IgG Secondary Antibody, Alexa Fluor 647 | Thermo Fisher Scientific | A-21245; RRID:AB_2535813 |
| Texas Red Streptavidin | Vector Laboratories | SA-5006, RRID:AB_2336754 |
| Sheep Anti-Digoxigenin Fab fragments Antibody, AP Conjugated, Roche | Roche | 11093274910; RRID:AB_514497 |
| **Chemicals, Peptides, and Recombinant Proteins** | | |
| Protein kinase Cζ pseudosubstrate, myristoyl trifluoroacetate salt | Sigma-Aldrich | P1614 |
| 1-Azakenpaullone | Sigma-Aldrich | A3734 |
| Biotinylated Dextran Amine-Texas Red | Vector Laboratories | SP-1140; RRID:AB_2336249 |
| Dextran, Alexa Fluor™ 488; 10,000 MW, Anionic, Fixable | Thermo Fisher Scientific | D22910 |
| Dextran, Alexa Fluor™ 555; 10,000 MW, Anionic, Fixable | Thermo Fisher Scientific | D34679 |
| Dextran, Alexa Fluor™ 647; 10,000 MW, Anionic, Fixable | Thermo Fisher Scientific | D22914 |
| Dextran, Cascade Blue™, 10,000 MW, Anionic, Lysine Fixable | Thermo Fisher Scientific | D1976 |
| **Deposited Data** | | |
| *Nematostella vectensis* genome assembly 1.0 | JGI | https://genome.jgi.doe.gov/Nemve1/Nemve1.home.html |
| **Experimental Models: Organisms/Strains** | | |
| *Nematostella vectensis* | Whitney Laboratory for Marine Bioscience, FL, USA. | RRID:SCR_005153 |
| **Oligonucleotides** | | |
| Genomic PCR *Nvpar-6* and dn*Nvpar-6*: F-AAAACCACCATCAGCCGAGTCA; R-TATTGATAGAATACCAGTCTCA |  | NEMVEDRAFT_v1g233358 |
| sgRNAs *Nvpar-6*: 1-GGATGTTGCCGACTCGCAGT; 2-GGAGAAGGCGAACTCGTCTG; 3-GGATAACCCTGTGCCAGTCA | CRISPRevolution sgRNA; Synthego | NEMVEDRAFT_v1g233358 |
| dn*Nvpar-3*: F-ATGATGAAGGTTGTAGT; R-TGCGCCCGATTCGAATCCATCT |  | NEMVEDRAFT_v1g240248 |
| sgRNAs *Nvpar-3*: 1-GGGTGTTCGAGGGACGCGAT; 2- GGGCAGGTTTATCCCGAAGG; 3- ACCAACGAUCUAGAUCCAGU | CRISPRevolution sgRNA; Synthego | NEMVEDRAFT_v1g240248 |
| Genomic PCR *Nvpar-3*: F-GTAGACGGGACTGGTTTGGA; R-AGGGACAGGTTGCTCCTTTT |  | NEMVEDRAFT_v1g240248 |
| dn*Nvpar-1*: F-AATATAAACTATGAACTTAACG; R-TTAAAGTTTTAATTCATTTGCA |  | NEMVEDRAFT_v1g 139527 |
| sgRNAs *Nvsnail-A*: 2-GGGGCCGGTAATGACGCGCG; 3-GGCGTAGAGTCACACCGCAA; 4-GGCGATGATATCGAGCTCGG; 5-GGGCATCTTGAGTGCACCCA | CRISPRevolution sgRNA; Synthego | NEMVEDRAFT_v1g240686 |
| sgRNAs *Nvsnail-A*: 1-GGGCTCTCTTGCTCCGTAAC; 6-GGGTTTCCTGGCGCTGGGAT | CRISPRevolution Modified sgRNA; Synthego | NEMVEDRAFT_v1g240686 |
| Genomic PCR *Nvsnail-A* (full length): F-ATGCCCCGCTCGTTTCTAG; R-TCCTTGTGACGGGCAGCC |  | NEMVEDRAFT_v1g240686 |
| sgRNAs *Nvsnail-B*: 1-GGAAGAGGATGTGAGGTTTT; 2-GAGATGATATTAGGCTGGTG; 4-GAAAAGCTGTACGACTCCTT; 5-GGGCATCTTGAGAGCGCCCA | CRISPRevolution sgRNA; Synthego | NEMVEDRAFT_v1g236363 |
| sgRNAs *Nvsnail-B*: 3-GGGTGAAGACTAAGACAGAG; 6-GGGCGATGAATCGTGTTTAA | CRISPRevolution Modified sgRNA; Synthego | NEMVEDRAFT_v1g236363 |
| Genomic PCR *Nvsnail-B* (full length): F-ATGCCGAGGTCCTTCCTGG; R-GCAGAGATTTTGCCGACACAT |  | NEMVEDRAFT_v1g236363 |
| **Recombinant DNA** | | |
| pSPE3-mVenus | Roure et al., 2007 |  |
| pSPE3-mCherry | Roure et al., 2007 |  |
| pSPE3*-Nvpar-6*-mVenus | Salinas-Saavedra et al., 2015 |  |
| pSPE3-*Nvpar-3*-mVenus | Salinas-Saavedra et al., 2015 |  |
| *Nv*ß-catenin expression constructs | Wikramanayake et al., 2003; Röttinger et al., 2012 |  |
| *Nvsnail-A* backbone to generate expression constructs | Magie et al., 2007 |  |
| **Software and Algorithms** | | |
| Fiji (ImageJ) | NIH | http://fiji.sc |
| Imaris 7.6.4 | Bitplane Inc. |  |
| CRISPRscan | Moreno-Mateos et al., 2015 | http://www.crisprscan.org/ |
