## Supplementary figures and images for "Germ layer specific regulation of cell polarity and adhesion gives insight into the evolution of mesoderm"

### Supplementary Materials

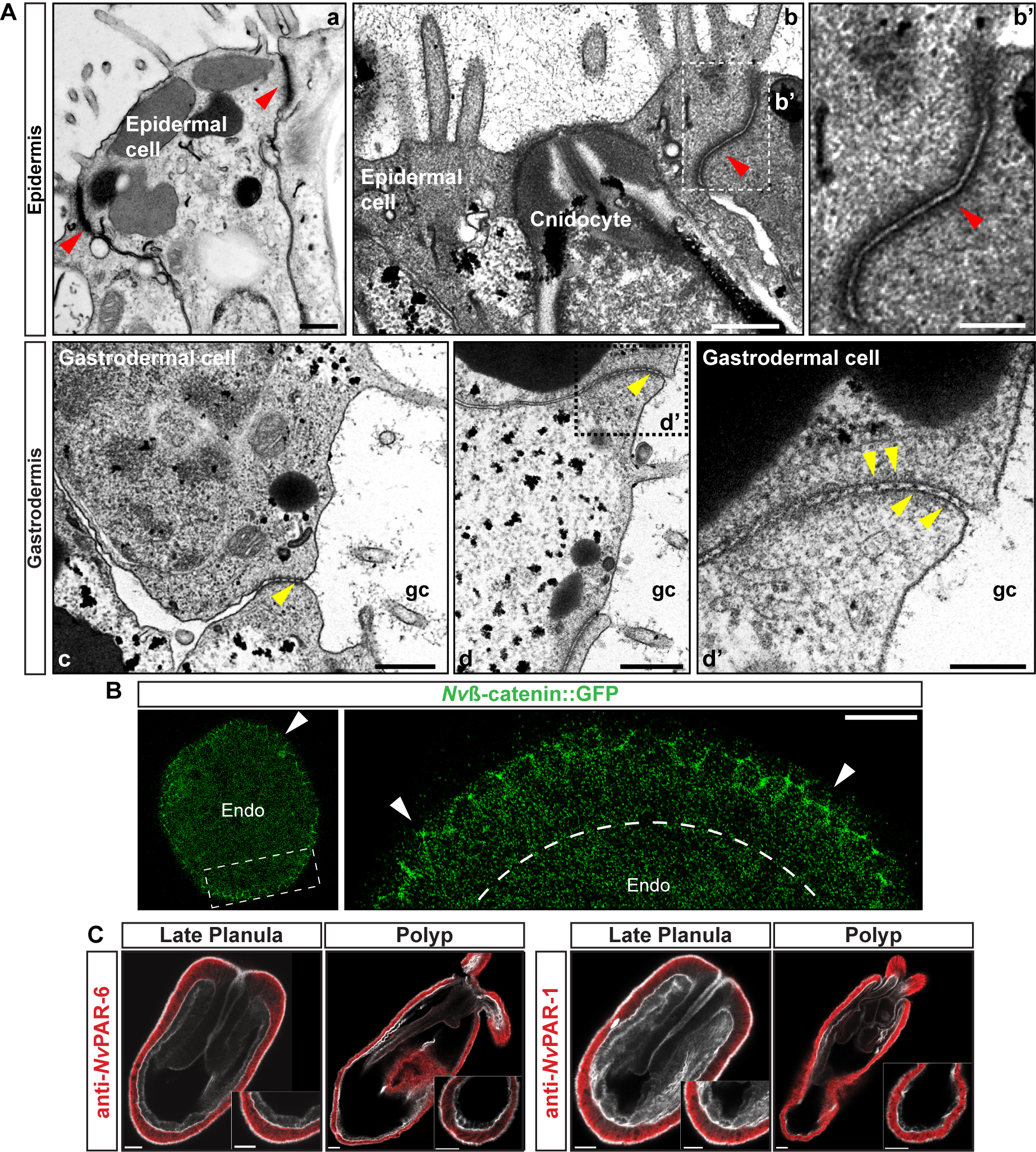

### Supplementary Materials

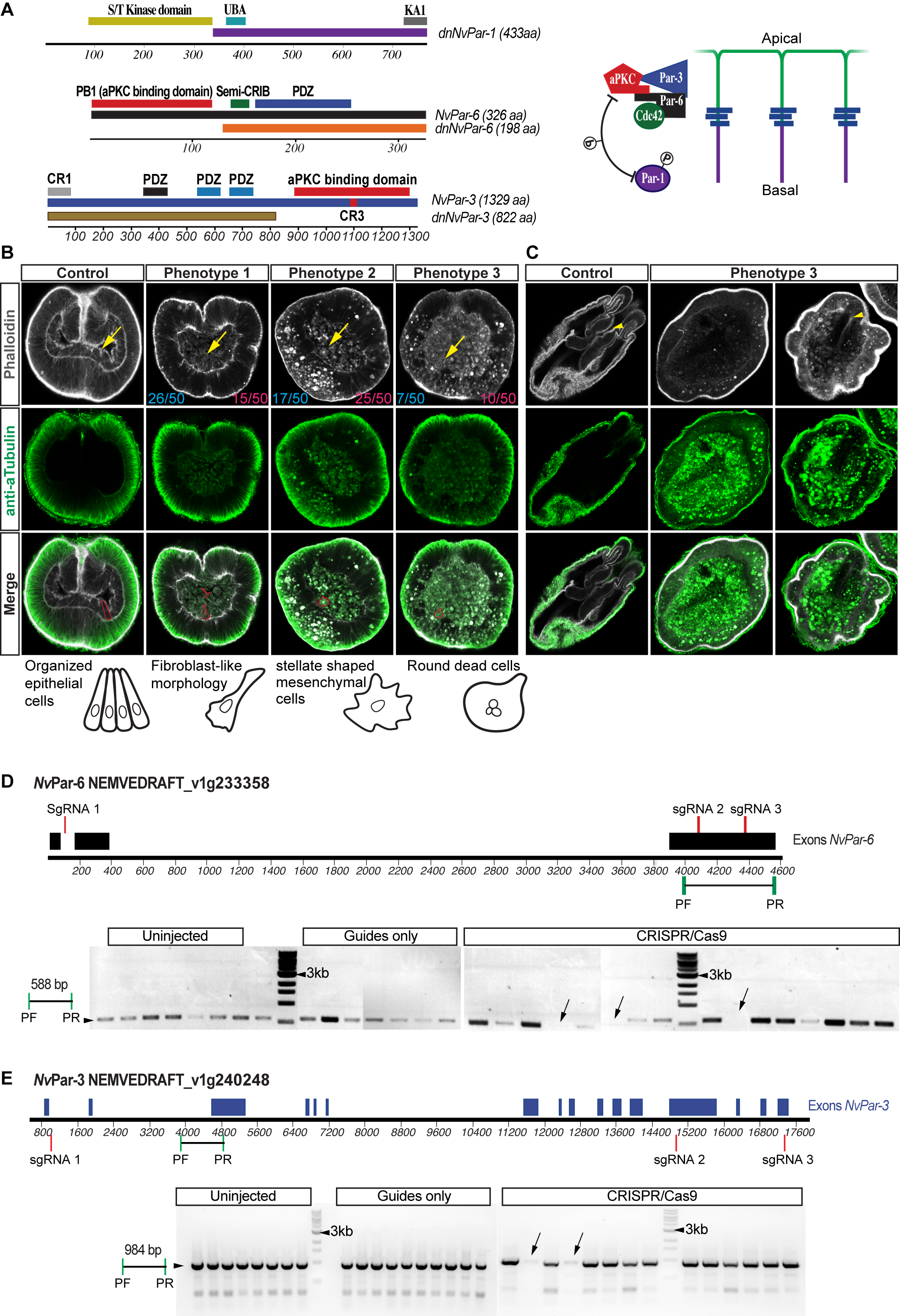

### Supplementary Materials

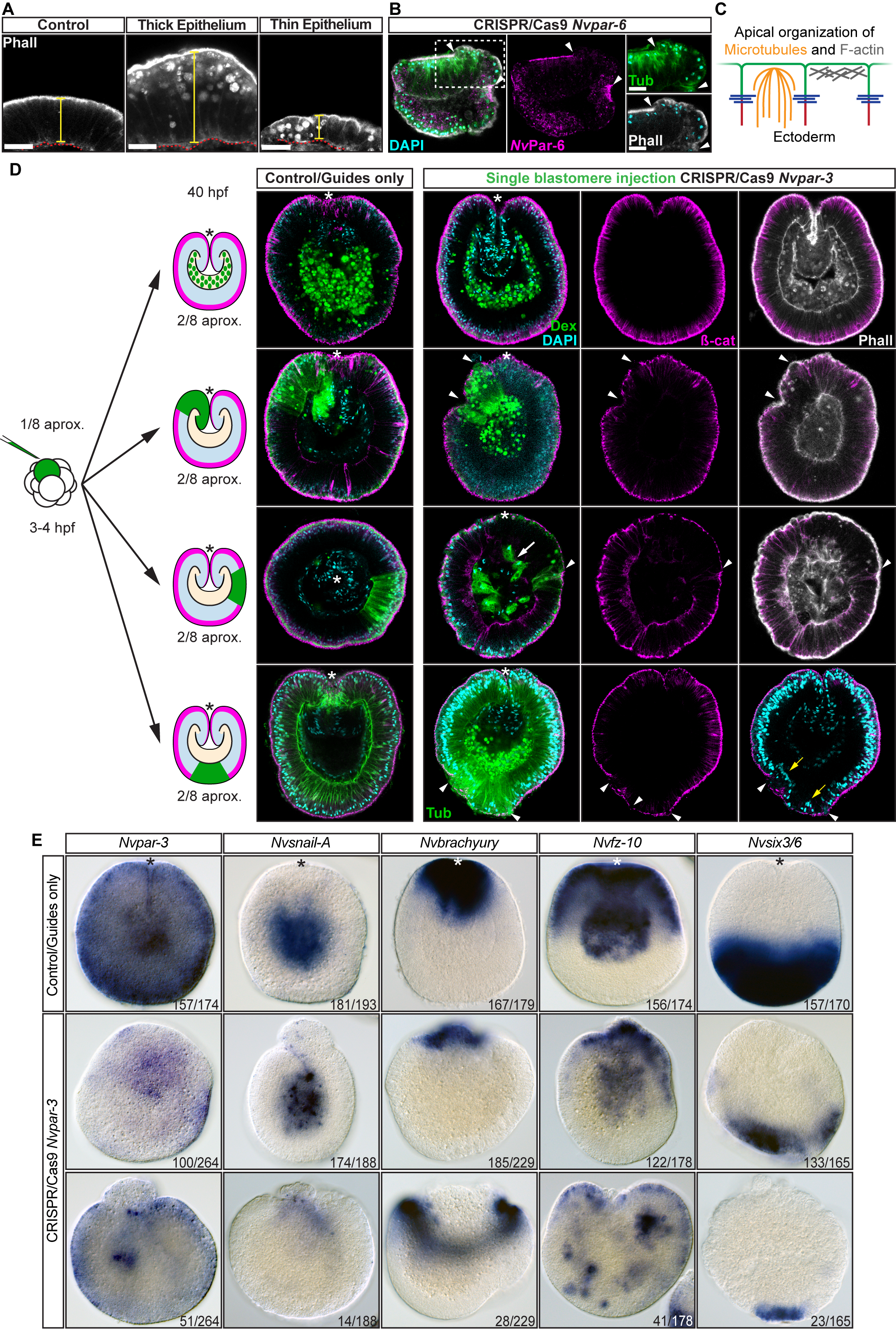

### Supplementary Materials

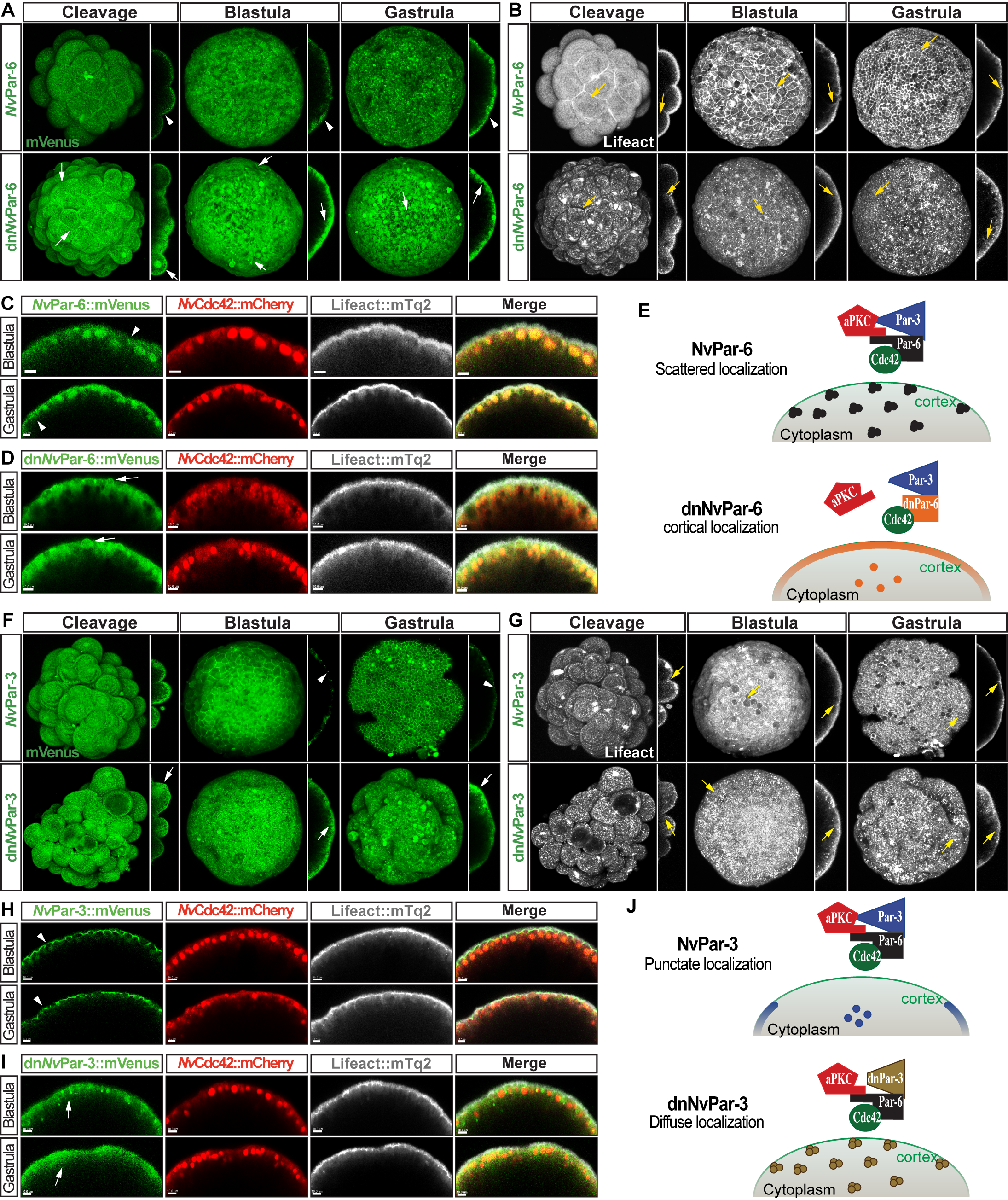

### Supplementary Materials

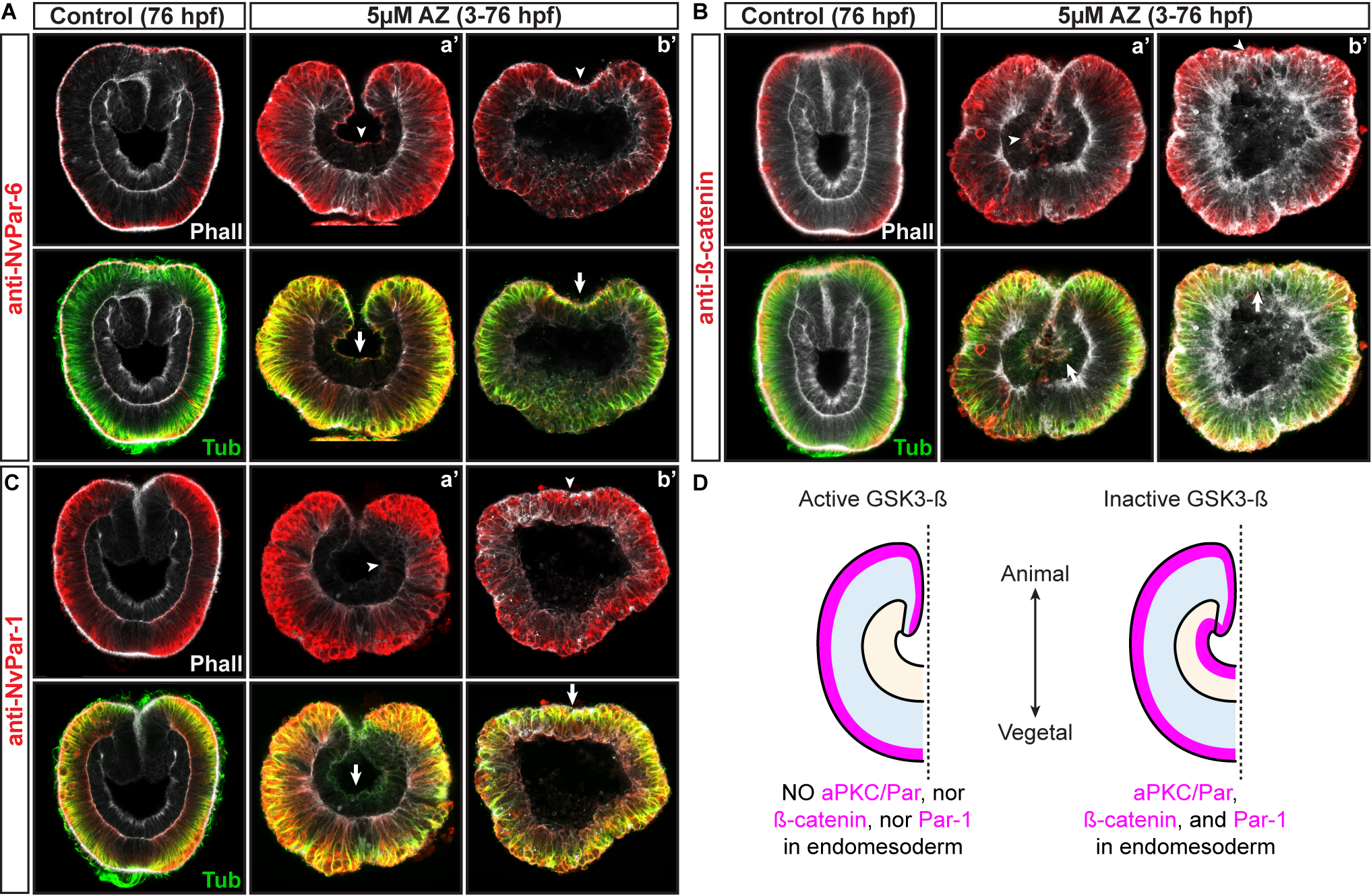

### Supplementary Materials

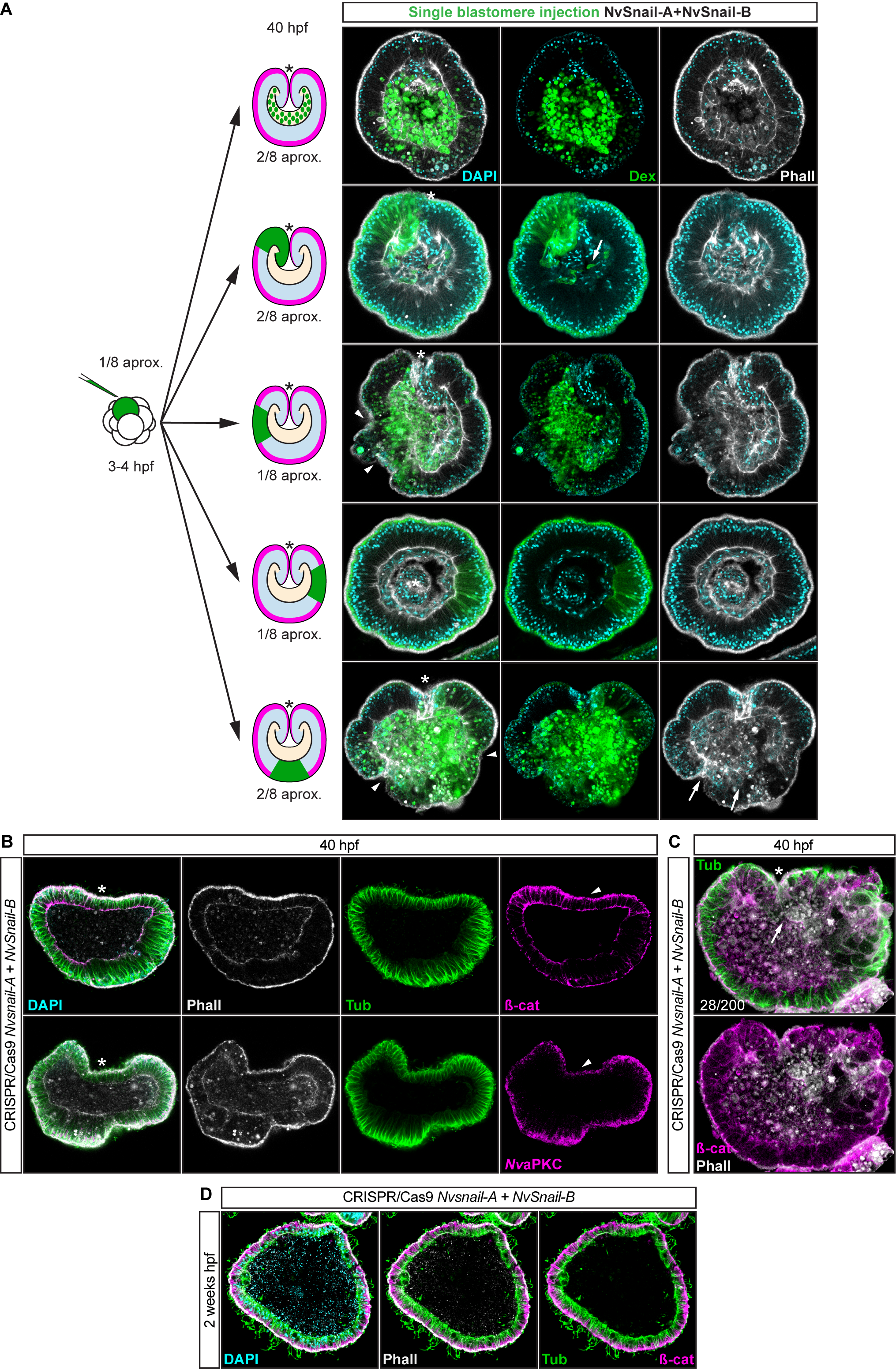

### Supplementary Materials

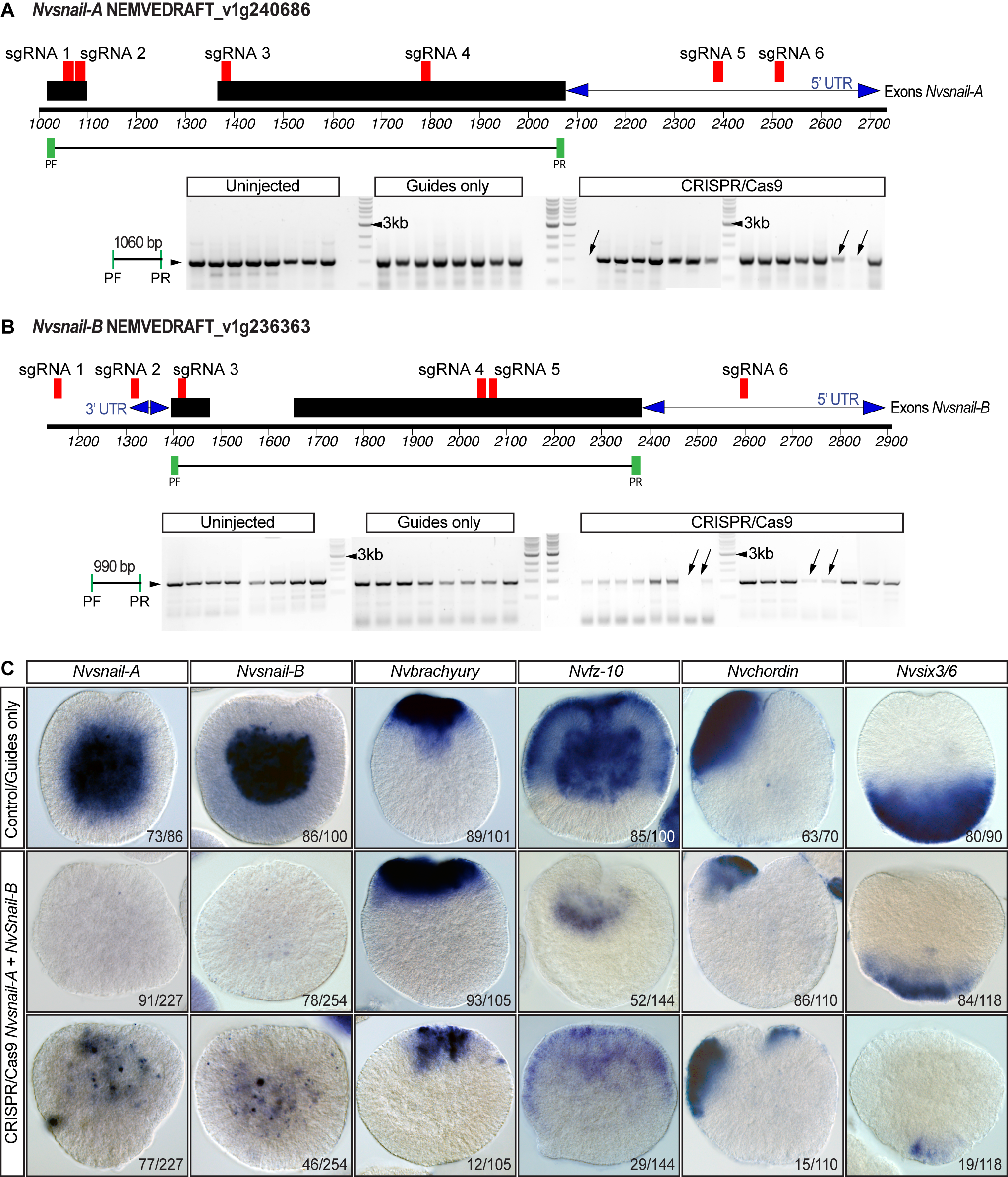

### Supplementary Materials

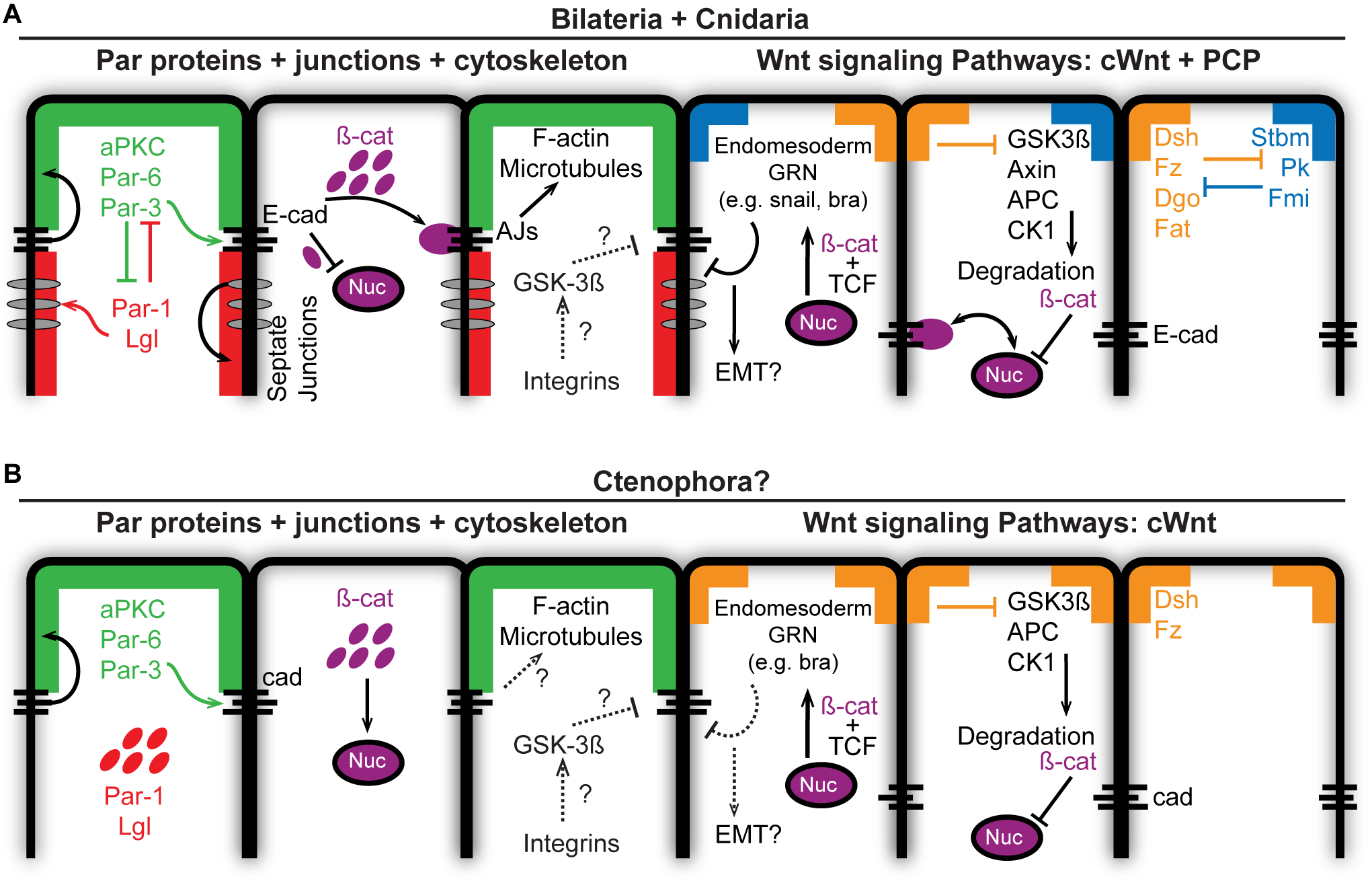
